# PHD3 controls energy homeostasis and exercise capacity

**DOI:** 10.1101/781765

**Authors:** Haejin Yoon, Jessica B. Spinelli, Elma Zaganjor, Samantha J. Wong, Natalie J. German, Elizabeth C. Randall, Afsah Dean, Allen Clermont, Joao A. Paulo, Daniel Garcia, Hao Li, Nathalie Y. R. Agar, Laurie J. Goodyear, Reuben J. Shaw, Steven P. Gygi, Johan Auwerx, Marcia C. Haigis

## Abstract

Rapid alterations in cellular metabolism allow tissues to maintain homeostasis during changes in energy availability. Acetyl-CoA carboxylase 2 (ACC2) provides a central regulation and is phosphorylated acutely by AMPK during cellular stress to relieve repression of fat oxidation. While ACC2 is hydroxlated by prolyl hydroxylase 3 (PHD3), the physiological consequence of PHD3 is little understood. We find that ACC2 phosphorylation and hydroxylation occur in a reciprocal fashion. ACC2 hydroxylation occurs in conditions of high energy and leads to decreased fatty acid oxidation. Furthermore, AMPK-mediated phosphorylation of ACC2 is inhibitory to hydroxylation. PHD3 null mice demonstrate loss of ACC2 hydroxylation in heart and skeletal muscle and display elevated fatty acid oxidation. Whole body or skeletal muscle-specific PHD3 loss enhanced exercise capacity during an endurance exercise challenge. In sum, these data identify PHD3 as a physiological regulator of energy balance during acute stress.

## INTRODUCTION

Metabolic adaptation plays a fundamental role in maintaining energy homeostasis. In many tissues, fatty acids are exploited as an adaptive fuel to enable survival under conditions of metabolic stress, such as nutrient deprivation or exercise (Galgani and Ravussin, 2008; Woods and Ramsay, 2011; Palm and Thompson, 2017; Efeyan et al., 2015). As one example of metabolic adaptation, cells utilize fatty acid oxidation (FAO) to respond to low energy conditions, cold temperature, oxidative stress, and exercise (Lee et al., 2016; Chouchani et al., 2016; Grunt, 2018; Herzig and Shaw, 2018). However, the mechanism for the dynamic regulation of fatty acid oxidation in response to metabolic stress is not completely understood.

A reduction in cellular ATP and increase in the AMP/ATP ratio leads to activation of AMP-activated protein kinase (AMPK) (Hardie et al., 2012; Gowans and Hardie, 2014). In response to high AMP/ATP, AMPK phosphorylates acetyl-CoA carborylase (ACC) 1/2. AMPK phosphorylation of ACC 1/2 inhibits its activity to convert acetyl-CoA into malonyl-CoA. In the cytosol, ACC1 generates pools of malonyl-CoA thought to be important for de novo lipogenesis, while ACC2 associated with the outer mitochondrial membrane generates malonyl-CoA to inhibit carnitine palmitoyltransferase 1 (CPT1), which mediates the first step of long chain fatty acid transport into the mitochondria. Thus, ACC2 phosphorylation by AMPK provides a readout of cellular energy/stress with direct ties to fat synthesis and oxidation.

ACC2 is also subject to positive regulation in the control of fatty acid metabolism via hydroxylation on P450 by proline hydroxylase domain protein 3 (PHD3) (German et al., 2016). Prolyl hydroxylase domain family members (PHDs, also called EGLN1-3) are alpha-ketoglutarate-dependent dioxygenases (Epstein et al., 2001). A previous study showed that tumors with low PHD3 exhibited decreased ACC2 hydroxylation and elevated fatty acid oxidation (German et al., 2016). By contrast, the role of PHD3 in the control of fat oxidation in energy homeostasis has not been examined fully.

Here, we show that PHD3 modifies ACC2 at proline 450 physiologically in response to nutrient and energy availability in cells and mouse tissues, and impacts lipid and energy homeostasis in whole body PHD3 KO mice and skeletal muscle-specific PHD3 KO mice. Intriguingly, AMPK-mediated phosphorylation of ACC2 inhibits PHD3 binding and hydroxylation of ACC2, demonstrating one mechanism for the reciprocal relationship between ACC2 hydroxylation and phosphorylation. Using whole body or muscle specific PHD3 null mice, we probe the physiological relevance of PHD3 during fasting and exercise challenges. PHD3 loss increases fatty acid oxidation and the loss of PHD3 is sufficient to increase exercise capacity. Together, these results suggest that PHD3 signaling may have potential to tune fatty acid oxidation in response to energy state.

## RESULTS

### ACC2 hydroxylation is sensitive to glucose and negatively regulated by AMPK

In this study we sought to examine a physiological role for PHD3 in the control of fatty acid oxidation. We previously reported that PHD3 physically interacts with ACC2, a central regulator of fat catabolism (German et al, 2016). While both PHD3 and ACC2 signaling potentially converge on ACC2 through distinct post-translational modifications (PTMs), whether these PTMs act in a concerted manner to regulate ACC2 activity has not been explored (Figure 1A). As PHD3 is an alpha-ketoglutarate-dependent dioxygenase, and activation of ACC2 would subsequently repress FAO, we reasoned that PHD3 may hydroxylate ACC2 in response to nutrient availability. To examine whether PHD3 activity was sensitive to fluctuations in nutrient availability, we probed ACC2 hydroxylation in mouse embryonic fibroblasts (MEFs) cultured in the presence of low (5 mM glucose) or high glucose (25 mM glucose) in the presence of 10% FBS (Figure 1B). Since AMPK is active by increased AMP/ATP conditions, we utilized low glucose culture media as a control (Salt et al., 1998; Laderoute et al., 2006). As expected, ACC2 phosphorylation was increased in the low glucose condition. By contrast, we observed a reciprocal hydroxylation in ACC2 (Figure 1B and 1C) as detected by reactivity with an antibody to proline hydroxylation. Importantly, the addition of 25 mM glucose caused ACC2 hydroxylation within 5 minutes, but not in cells treated with the pan-PHD inhibitor dimethyloxallyl glycine (DMOG) (Figure 1B and 1C). DMOG treatment significantly decreased hydroxyl ACC2 levels compared to untreated control cells in high glucose media (Figure 1B and 1C). Finally, we tested whether other energetic inputs affected ACC2 hydroxylation (Figure S1A). In contrast to glucose, the addition of amino acids, fatty acids or dialyzed serum resulted in little to no induction of ACC2 hydroxylation (Figure 1D, Figure S1B-C). In sum, these data suggest a rapid induction in ACC2 hydroxylation in response to glucose abundance.

**Figure 1.**
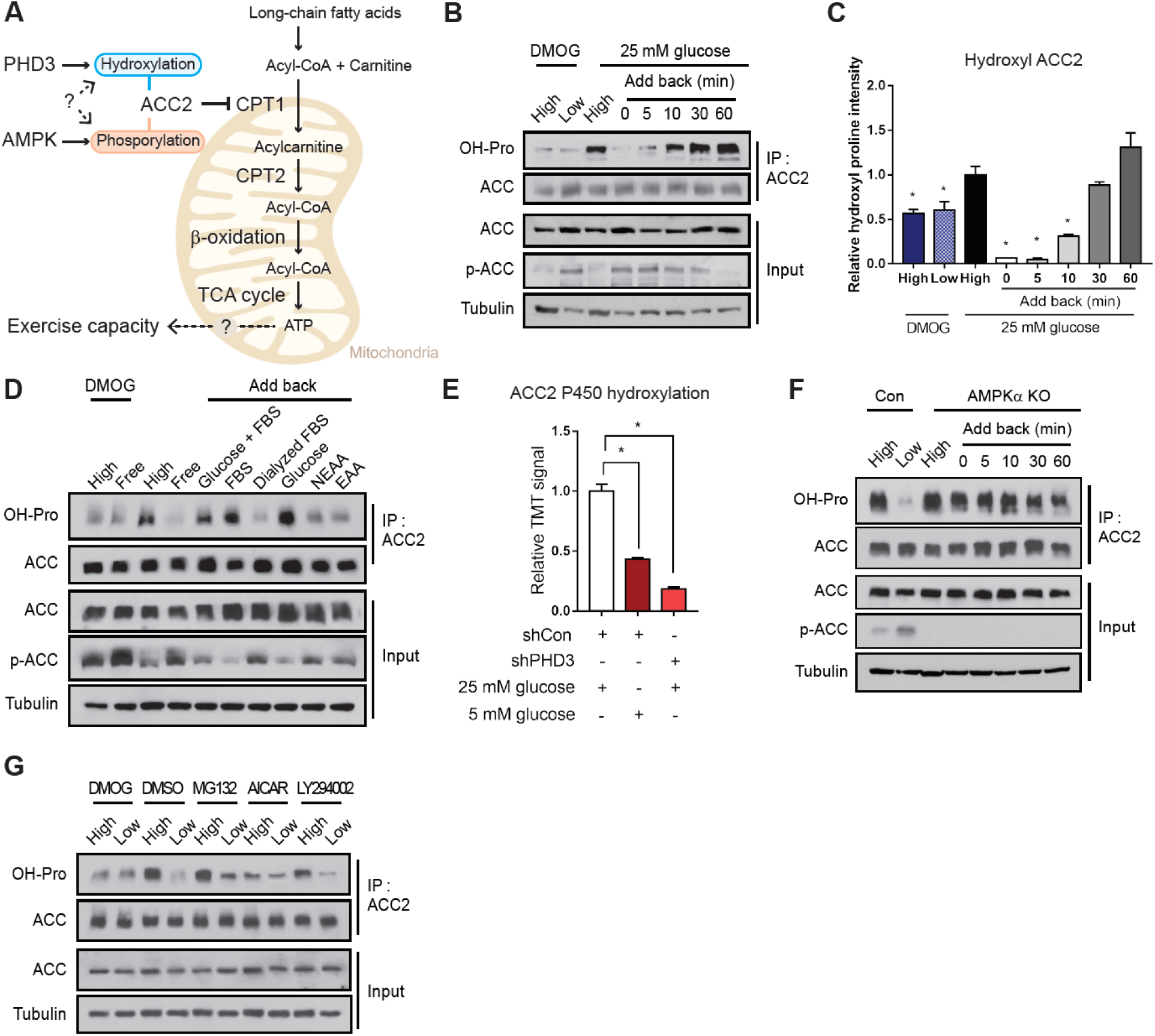
ACC2 hydroxylation is energy-sensitive and negatively regulated by AMPK. (A) Schematic of fates of post-translational modification on ACC2. (B) Immunoblots of anti-ACC2 immuno-precipates and corresponding whole-cell lysates of MEFs. MEFs were grown either in high (25 mM) or low (5 mM) glucose media for 12 hr with or without PHD inhibitor DMOG (1 mM). (C) Quantitation of western blotting of (B). Hydroxyl ACC2 was normalized to total bound ACC levels. The ratio of hydroxyl ACC2: total ACC2 was normalized to respective tubulin level (n=3). (D) ACC2 hydroxylation in MEFs in high versus low glucose, and in the indicated fuel. DMOG, 1 mM Dimethyloxallyl glycine; High, 25 mM glucose with serum; -Glu/-FBS, no glucose without serum; Glucose+FBS, 25 mM glucose with serum; FBS, no glucose with serum;Dialyzed FBS, no glucose with dialyzed FBS; Glucose, 25 mM glucose without serum; NEAA, no glucose with 1% non essential amino acids; EAA, no glucose with 1% essential amino acid. (E) Representative TMT signals identifying hydroxylated P450 on ACC2 in 293T cells in control (white), 5 mM glucose media (dark red) or PHD3 knockdown cells (red). (F) Immunoblots of ACC2 hydroxylation from control or AMPKa knock out MEFs in high versus low glucose, and upon 25 mM glucose re-addition (0, 5, 10, 30 and 60 min). (G) ACC2 hydroxylation in MEFs in high or low glucose, in the presence of DMOG (1 mM for 1 h), DMSO, MG132 (10 mM for 1 h), AICAR (1 mM for 1 h) and LY294002 (50 mM for 1 h).

We utilized Tandem Mass Tag (TMT) mass spectrometry to quantify changes in ACC2 proline modifications in response to glucose or PHD3 (Paulo, 2016; Navarrete-Perea et al., 2018; Berberich et al., 2018). In ACC2 immunoprecipitates, we observed a number of proline residues modified with a +15.9949 Th shift (Figure S1D), consistent with modification by hydroxylation. However, only P450 demonstrated elevated signal of a +15.9949 Th shift in a glucose- or PHD3-sensitive manner. The P450 +15.9949 Th shift was diminished 5.3-fold in PHD3 KD cells compared with control cells, and the P450 15.9949 Th shift was reduced 2.3-fold in wildtype control cells cultured in low glucose compared with high glucose media (Figure 1E, Figure S1E-G). These data are similar in magnitude to the fold-changes observed in ACC2 hydroxylation detected by Western blotting (Figure S1H).

Of interest, our studies suggested that ACC2 was phosphorylated in a reciprocal manner to its proline modification (Figure 1D). This observation suggested some level of circuitry between AMPK and PHD3 activity, and so we aimed to understand whether AMPK influenced PHD3 activity or vise versa. First, we tested whether PHD3 activity was affected by AMPK loss of function by measuring ACC2 hydroxylation in MEFs lacking the alpha catalytic subunit of AMPK (Figure 1F). Strikingly, the loss of AMPKα led to constitutive hydroxylation of ACC2 in a manner that was insensitive to glucose availability (Figure 1F). Cells treated with 5-aminoimidazole-4-carboxamide ribonucleotide (AICAR), which leads to constitutive activation of AMPK, decreased ACC2 hydroxylation, suggesting that AMPK-mediated phosphorylation represses ACC2 hydroxylation (Figure S1I). PHD3 expression was unchanged by ACC2 depletion or energy stress (Figure 1B, Figure S1J-L), and the depletion of PHD3 in MEFs did not alter ACC2 protein or mRNA level. We also tested whether AMPK activity was regulated by PHD3 and found that PHD3 KD did not affect the levels or kinetics of ACC2 phosphorylation.

We more broadly investigated whether modulation of other energy or nutrient sensing signaling pathways affected ACC hydroxylation. PI3K/AKT and mTOR signaling are known to coordinate with AMPK signaling (Manning and Toker, 2017; Inoki at el., 2003). Cells treated with LY294002 (PI3K/AKT inhibitor) or MG132 (proteasome inhibitor) did not significantly alter ACC2 hydroxylation status compared to the DMSO control (Figure 1G), while AICAR treatment decreased the reactivity of ACC2 with a pan-hydroxyl antibody (Figure 1G). These data suggest that AMPK-mediated phosphorylation of ACC2 is upstream and inhibitory to PHD3-mediated hydroxylation.

We then examined whether PHD3 and ACC2 hydroxylation were sensitive to changes in nutrients *in vivo*. ACC2 loss results in increased overall energy expenditure in mice, demonstrating a key role for fatty acid metabolism in physiological energy homeostasis (Choi et al., 2007). In order to activate AMPK and energy stress *in vivo*, we fasted mice for 16 hours and immunoprecipitated ACC2 from heart and quadriceps muscle to assess hydroxylation and phosphorylation. These tissues were probed because they expressed the highest levels of PHD3 compared to brain, lung, liver, kidney, spleen, testis and white adipose (Figure 2A). Although PHD3 expression was unchanged with fasting (Figure 2B), ACC2 reactivity with an antibody to proline hydroxylation was significantly decreased in heart and quadriceps from fasted mice relative to fed mice (Figure 2C). Conversely, phosphorylated ACC2 was increased after fasting in both tissues, consistent with elevated AMPK activity under these conditions (Figure 2C).

**Figure 2.**
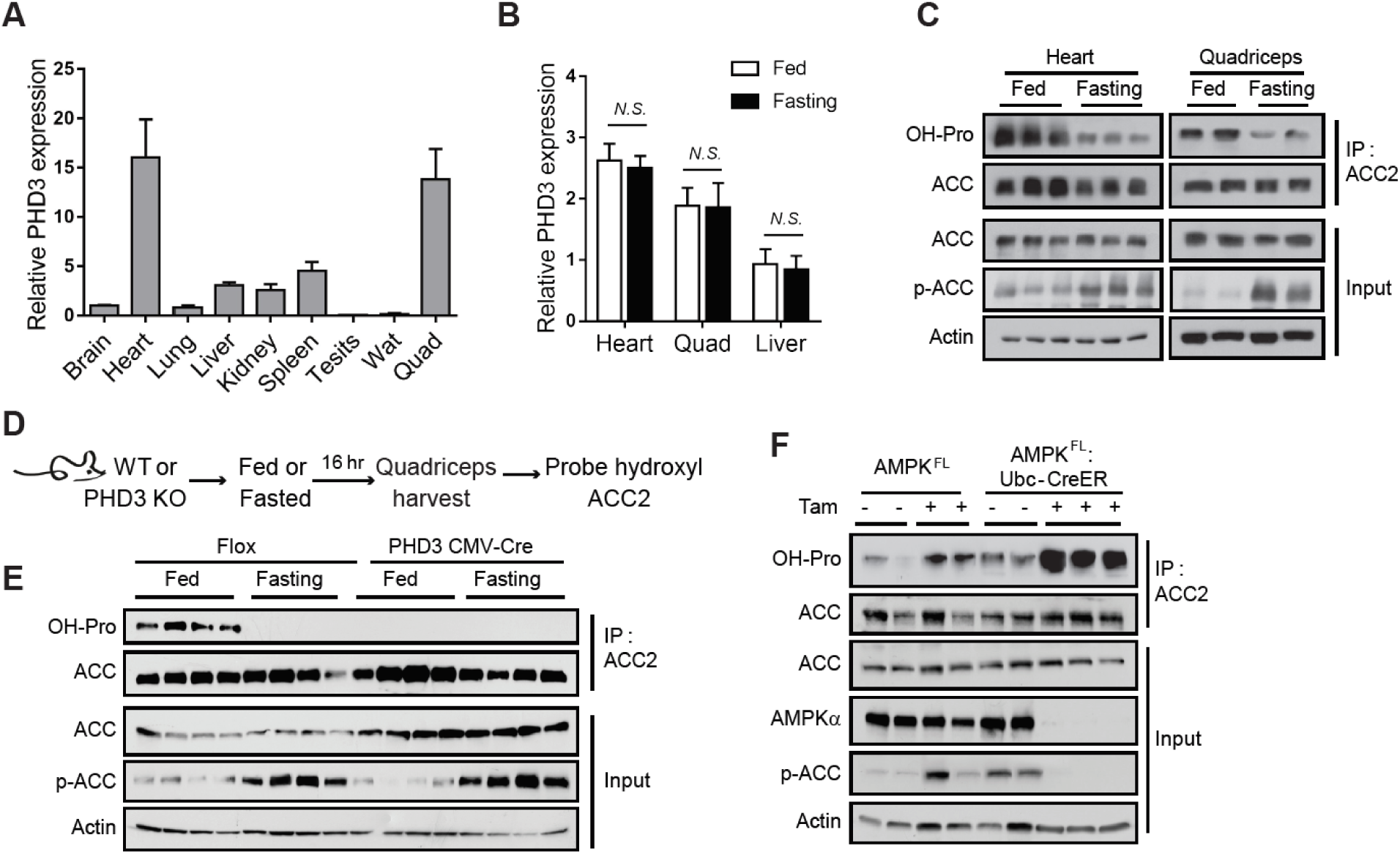
PHD3-mediated ACC2 hydroxylation in mouse tissues. (A) PHD3 mRNA levels in mouse brain, heart, lung, liver, kidney, spleen, testis, white adipose tissue, and quadriceps (n = 4). (B) PHD3 mRNA levels in mouse heart, quad and liver under fed or fasting conditions (n = 4). (C) Immunoprecipitation (IP) and western blot analysis of ACC hydroxylation from whole-cell lysates of mouse tissues that were fasted or fed for overnight. (D) Schematic of the detection of ACC2 hydroxylation in WT or PHD3 KO mouse quadriceps under fed or fasted condition. (E) Immunoprecipitation and western blot analysis of ACC2 hydroxylation and phosphorylation in WT or PHD3 KO mouse quadricep muscles in fed and fasted conditions, N = 4. (F) Immunoprecipitation and western blot analysis of ACC2 hydroxylation in quadriceps from AMPK-WT and AMPK-KO mice. AMPKFL;Ubc-CreER mice were treated with tamoxifen (4mg per day, for 5 days) to induce knockout of AMPK. Quadriceps were harvested 3 weeks post-tamoxifen treatment and protein was extracted.

To determine if PHD3 was required for ACC2 hydroxylation *in vivo*, we examined tissues from PHD3 knockout (KO) mice (Figure S2A-C) (Takeda et al., 2006). PHD3 was requisite for the hydroxylation of ACC2 in quadriceps muscle, as measured by cross-reactivity using an antibody for proline hydroxylation (Figure 2D and 2E). Loss of PHD3 did not affect ACC2 phosphorylation, consistent with cellular data that PHD3 activity does not act upstream of AMPK (Figure 2E). To determine if AMPK regulates ACC2 hydroxylation *in vivo*, we measured ACC2 hydroxylation in AMPKα KO mice. AMPK^FL^-Ubc-Cre^ER^ (AMPKα KO) mice were treated with 4 mg of tamoxifen for 5 days to knockout AMPK, and ACC2 hydroxylation was probed in AMPKα KO mice quadriceps using a antibody reactive to pan-proline hydroxylation (Figure 2F). As expected, ACC2 was not phosphorylated in these animals due to loss of AMPK catalytic activity (Figure 2F). Consistent will cellular data, loss of AMPK activity *in vivo* increased ACC2 hydroxylation signal compared to control mice (Figure 2F). Taken together, these findings demonstrate that the ACC2 hydroxylation signal is dependent on PHD3 in vivo, sensitive to the energy status of tissues, and repressed by AMPK.

### ACC2 and PHD3 binding is interrupted by serine 222 phosphorylation of ACC2

As ACC2 has been shown to physically bind with PHD3 (Tran et al., 2015; German et al., 2016), we sought to elucidate whether this binding interaction was sensitive to cellular nutrient status. ACC2 is a 280-kDa multi-domain enzyme, comprised of: N-terminal (NT), biotin carboxylase (BC), carboxyltransferase (CT), and biotin carboxyl carrier protein (BCCP) (Figure 3A, Figure S3A-B) (Tran et al., 2015; Tong et al., 2006). ACC2 residues S222 and P450 reside in the NT and BC domains, respectively (Figure 3A, Figure S3A), which are modeled to form a dimer with an interface adjacent to S222 phosphorylation sites, whereas P450 resides on the outside of the dimer interface (Figure 3A). We engineered 293T cells to express C-terminal-Flag-tagged ACC2 and C-terminal-HA-tagged PHD3. WT-PHD3 and WT-ACC2 co-immunoprecipitated to a greater extent in cells cultured in high glucose compared to low glucose (Figure 3B). As AMPK reduced PHD3 activity, we tested whether ACC phosphorylation interfered with PHD3 binding by measuring the co-immunoprecipitation of PHD3-HA with WT ACC2, S222A ACC2, or S222E ACC2. PHD3 co-immunoprecipitated with an S222A variant of ACC2 with greater intensity compared with WT ACC2, suggesting that phosphorylation may be inhibitory to this binding interaction (Figure 3C).

**Figure 3.**
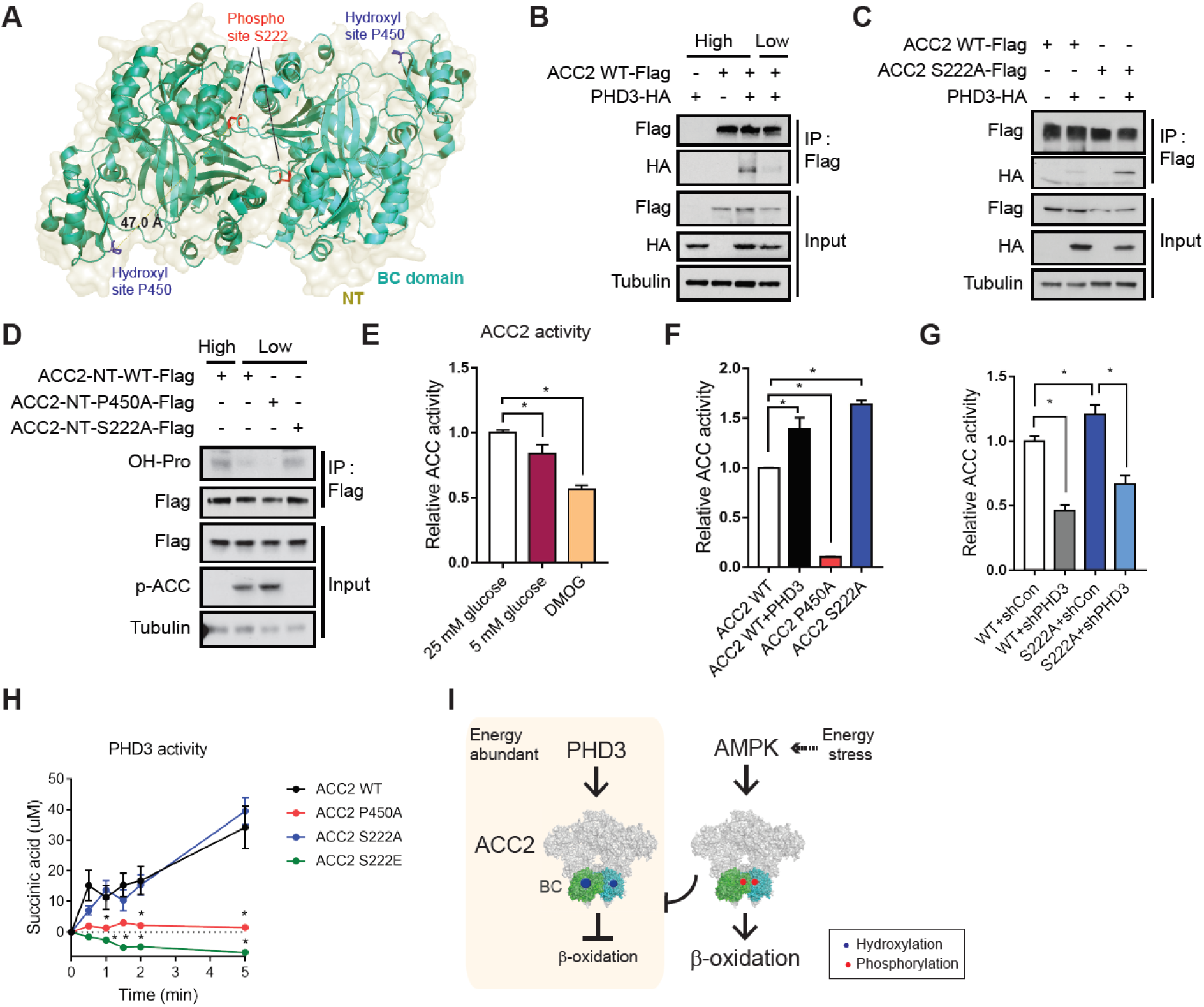
ACC2 S222 phosphorylation inhibits ACC2 hydroxylation and activity. (A) P450 (hydroxylation site, blue) is located m the ACC2 BC domain (green and cyan PDB: 3JRW). The distance between S222 (phosphorylation site, red) and hydroxylation sites is approximately 47 A. (B) HEK293T cells were transfected with ACC2-Flag and/or PHD3-HA. Cells were incubated with 25 mM high glucose media or 5 mM low nutrient media tor 8 h. Proteins in cell lysates were precipitated by anti-Flag antibody and lmmuno-blotted using the indicated antibodies. (C) IP and western blot analysis ot PHD3 and ACC2 interaction in the presence or absence of S222A mutation on ACC2. (D) Phosphorylation of ACC2 interrupts the binding between ACC2 and PHD3. ACC2 N-terminus (NT)-WT-Flag, ACC2 NT-P450A-Flag, ACC2 NT-S222A-Flag and/or PHD3-HA were transfected into HEK293T cells. Proteins were immunoblotted using the indicated antibodies. (E) ACC activity assay m HEK293T cell lines in 25 mM glucose (high nutrient), 5 mM glucose (low nutrient), or DMOG. Activity was normalized to protein level (n=4). (F) ACC activity assay m MEFs overexpressing WT ACC2, ACC2 P450A, ACC2 S222A or PHD3 (n = 3). (G) ACC activity assay m MEFs overexpressing WT ACC2, ACC2 P450A, ACC2 S222A, or PHD3 (n-4). (H) In vitro hydroxylation assay for PHD3 enzymatic activity with recombinant protein about N-terminus domains of ACC2 WT, ACC2 P450A, S222A, or S222E, and PHD3 (n=3). (I) Schematic figure that AMPK-induced phosphorylation inhibits PHD3-related hydroxylation on ACC2 dimer structure. Posttranslational modifications by AMPK (red dots) and PHD3 (blue dots) on the ACC2 BC domain are indicated m the estimated ACC2 model (PDB: 5CSL).

To begin to biochemically define the interaction between these proteins, we mapped the domain(s) on ACC2 important for binding to PHD3 (Figure S3C-H). Each domain was expressed recombinantly in *E. coli* with a C-terminal GST-tag, purified, and incubated *in vitro* with recombinant PHD3-HA-His to test their ability to co-immunoprecipitate. We observed that the BC domain co-immunoprecipitated with PHD3 *in vitro*, while the CT and BCCP domains demonstrated no interaction (Figure S3C-E). The reciprocal immunoprecipitation with anti-HA (Figure S3F-H) also identified selective binding between PHD3 and the BC domain (Figure S3C-I).

Since AMPK activity represses ACC2 hydroxyl modification, we tested whether S222 phosphorylation blocks the PHD3-ACC2 binding interaction *in vitro*. We generated expression construct to overexpress the N-terminal domain of WT ACC2 containing amino acids 1-763 (ACC2-NT-WT) or an AMPK phospho-mutant (ACC2-NT-S222A) in 293T cells (Figure S3B). This truncated variant contained the biotin carboxylase and ATP grasp domains but migrates distinctly from full length ACC (∼250 kDa), allowing us to monitor the effect of each PTM. We transfected WT ACC2, P450A ACC2 or S222A ACC2 into 293T cells and measured hydroxylation in immunoprecipitated ACC2 (Figure 3D). ACC2-NT-WT co-immunoprecipitated with PHD3 in cells and was hydroxylated (Figure 3D, Figure S3I-J). ACC2 hydroxylation was decreased in low nutrient compared to high nutrient condition in ACC2-NT-WT. In low glucose, ACC2-NT-P450 demonstrated lower hydroxylation (Figure 3D), while the constitutively dephosphorylated mutant ACC2-NT-S222A was more hydroxylated.

We next examined the effect of glucose, PHD3, and ACC2 modifications on ACC2 activity in a direct, biochemical assay of ACC2 activity using radioactive sodium bicarbonate on immunoprecipitated ACC2. ACC2 activity was decreased in 5 mM glucose and DMOG treatment compared to WT cells in 25 mM glucose (Figure 3E). To test if ACC2 activity was affected by PHD3, we measured ACC2 activity assay of WT ACC2, P450A ACC2 or S222A ACC2 in PHD3 overexpressing 293T cells (Figure 3F). ACC2 activity increased in the presence of PHD3. P450A ACC2 demonstrated low activity, while S222A ACC2 demonstrated increased activity compared with WT ACC2 (Figure 3F). Furthermore, ACC2 activity was reduced by PHD3 knockdown in both conditions (Figure 3G).

To probe if the phosphorylation on S222 affects the ability of ACC2 to be hydroxylated we measured the activity of purified, recombinant PHD3 using the N-terminal domain of ACC2, P450A ACC2, S222A ACC2, or S222E ACC2 as substrates. PHD3 uses oxygen and α-ketoglutarate to hydroxylate target proline residues, generating carbon dioxide and succinate in the process. Thus, to quantify PHD3 activity, we measured the production of succinate over the course of the reaction (Figure 3H). While PHD3 demonstrated activity using WT ACC2 as a substrate, PHD3 possessed no detectible activity towards a control ACC2 P450A mutant. PHD3 demonstrated significant activity using the S222A ACC2 variant as a substrate; however, the phospho-mimetic S222E ACC2 variant was a poor substrate of PHD3. These data are consistent with the model that phosphorylation of ACC2 at S222 is inhibitory to PHD3-mediated hydroxylation of ACC2 (Figure 3I).

### PHD3 represses fat catabolism *in vivo*

As long chain fatty acids are prepared for the multi-step process of mitochondrial import, they must be converted to acyl-carnitines by CPT1, which is inhibited by ACC2 (McCoin et al., 2015) (Figure 4A). Thus, we assayed the effect of ACC2 PTMs on palmitate oxidation. In MEFs, low glucose conditions increased FAO, and was further exacerbated by PHD3 reduction (Figure 4B). We repeated these experiments in C2C12 cells, which are mesenchymal stem cells classically used as a model of muscle cells (Millay et al., 2013). Similar to HEK293Ts and MEFs, fatty acid oxidation was increased in C2C12 cells with reduced PHD3 (Figure S4A). Moreover, PHD3 knockdown repressed ACC2 hydroxylation in C2C12 cells (Figure S4B).

**Figure 4.**
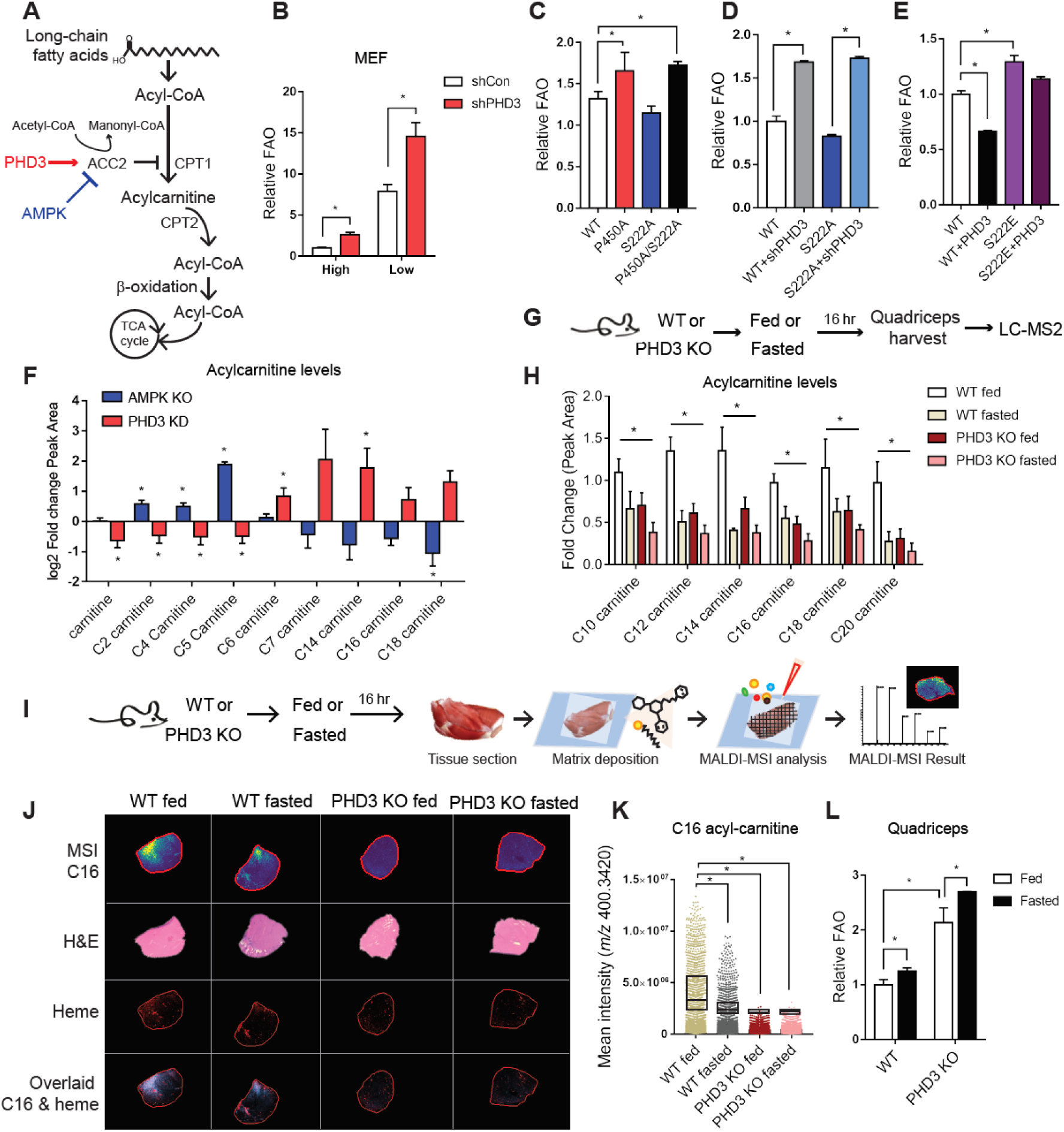
Loss of PHD3 results in increased fatty acid catabolism in mouse muscle. (A) Schematic of the regulation of long chain fatty acid oxidation by PHD3 and AMPK. (IB) Palmitate oxidation in WT or PHD3 knockdown MEFs under high or low nutrient condition using FAO analysis (n = 3). (C) Palmitate oxidation assay in MEFs with overexpressed ACC2 WT, ACC2 P450A, ACC2 S222A or ACC2 P450A/S222A double mutant (n = 3) using FAO analysis. (D) FAO analysis in MEFs overexpressed ACC2 WT or ACC2 S222A with PHD3 silencing (n = 3). (E) Palmitate oxidation assay in MEF cell lines with overexpression ACC2 WT, S222E, or PHD3 (n=3). (F) Relative abundance of acylcamitines in MEFs depleted of PHD3 (red) and AMPK (blue) compared to control cells. For all comparisons two-tailed t test was used. N = 4, P < 0.05. (G) Flowchart of the metabolomic analysis using LC-MS in WT or PHD3 KO mouse under fed or fasted condition. (H) The relative metabolite abundance of long chain acyl-carnitines in WT or PHD3 KO mouse quadriceps under fed or fasted condition (n = 4). (I) Flowchart of the metabolomics analysis in WT or PHD3 KO mouse quadriceps under fed or fasted condition. (J) Matrix-assisted laser desorption/ionization mass spectrometry imaging (MALDI-MSI) of palmitoylcamitine (Cl 6 acyl-carnitine, m/z 400.3420, Appm = 1.16) in WT or PHD3 KO mouse quadriceps under fed or fasted conditions, alongside H&E stained serial sections. Comparison of distribution of different ions – acyl-carnitine distribution correlates with vasculature. (K) The scatter dot plot of relative abundance of palmitoyl-carnitine under each condition. (L) Palmitate oxidation in WT or PHD3 KO mice quadriceps under fed or fasted condition using 14C-labeled palmitate and CO2 capture analysis (n=4).

We next examined whether AMPK affected the ability of PHD3 to regulate FAO. AMPK-mediated phosphorylation of ACC2 activates FAO (Cho et al., 2010; O’Neill et al., 2014). In high glucose, palmitate oxidation was not significantly altered in MEFs overexpressing the S222A ACC2 mutation compared to WT ACC2 (blue bar) cells cultured in high glucose (Figure 4C). As expected, MEFs overexpressing the P450A mutation displayed elevated palmitate oxidation compared with WT MEFs (Figure 4C). Cells expressing P450A/S222A ACC2 had increased FAO compared to WT (20.7%), similar to the P450A mutation alone (Figure 4C). We also measured FAO in the presence or absence of PHD3 knockdown and S222A ACC2. When PHD3 levels were reduced, FAO was increased in both ACC2 WT or S222A ACC2 expressing cells (Figure 4D). Conversely, PHD3 overexpression repressed fat oxidation (Figure 4E). Finally, based on our observation that AMPK activity repressed ACC hydroxylation, we tested whether the expression of S222E ACC2 would restore fat oxidation in the presence of PHD3 overexpression. Indeed, fat oxidation remained elevated in cells expressing both S222E ACC2 and PHD3 (Figure 4E).

As an alternate reporter of fat oxidation, we monitored acyl-carnitine and metabolite profiles (Figure S4C-E). Compared with respective controls, PHD3 KD MEFs possessed elevated long chain acyl-carnitines, while AMPK KO MEFs showed reduced acyl-carnitines with an inflection point at the junction of short versus long chain acyl-carnitines (Figure S4C and Figure 4F). Of note, the level of ACC2 or mitochondrial mass as measured by MitoTracker was unchanged by PHD3 loss (Figure S4F-J). Since glucose and fatty acids are both converted to acetyl-CoA, activated fat oxidation may suppress glycolysis (De Leiris et al., 1975; Randle et al., 1963; Jenkins et al., 2011). To examine whether PHD3 regulates glucose oxidation in addition to fat oxidation, we measured the basal oxygen consumption rate (OCR) resulting from glucose or fat oxidation WT or PHD3 knockdown MEFs using a Seahorse flux analyzer (Figure S4K-L). Basal glucose-driven respiration was unchanged in shPHD3 MEFs under high or low nutrient conditions compared to WT cells (Figure S4L). By contrast, loss of PHD3 increased basal fat oxidation-driven respiratory activity, consistent with increased FAO (Figure S4K).

To identify the changes in lipid metabolism driven by PHD3 *in vivo*, we analyzed the levels of long chain acyl-carnitines in WT or PHD3 KO mice quadriceps using LC-MS/MS (Figure 4G). C12, C14, C16 and C18 acyl-carnitine levels were decreased in WT quadriceps under fasted conditions in PHD3 KO mice compared to WT mice (Figure 4H). In quadriceps we observed long chain acyl-carnitines were lower in WT mice under fasted conditions compared to fed WT mice (Figure 4H). These long chain acyl-carnitines were even lower in PHD3 KO relative to WT mice and remained low under fed or fasted conditions. Our findings were corroborated using matrix-assisted laser desorption/ionization mass spectrometry imaging (MALDI-MSI) to assess spatial resolution of metabolite profiles *in vivo* (Figure 4J). Consistent with LC-MS/MS data, acyl-carnitines (C14 acyl-carnitine, C16 acyl-carnitine, and C18 acyl-carnitine) levels were lower in the quadriceps of PHD3 KO mice under fed conditions compared to those in WT mice in fed conditions, whereas there was no further difference in fasted PHD3 KO mice (Figure 4J-K, S5M-N). Heme levels were used for reference, which were not changed in PHD3 KO tissue (Figure 4J, S5O). Interestingly, we observed that while low glucose conditions in cells increased acyl-carnitine levels, these conditions of fasting in mice decreased C14, C16 and C18 acyl-carnitines. One account for this difference could be that these fuels might have a higher oxidization rate during fasting *in vivo*, which would prevent the acyl-carnitines from accumulating. To test this idea, we measured the rate of ^14^C-palmitate oxidation (Figure 4L) on quadriceps slices isolated from WT and PHD3 KO mice. Palmitate oxidation increased in PHD3 KO mice quadriceps compared to WT tissue (Figure 4L) and increased in both groups after a 16 hour fast (Figure 4L). These alterations appeared in the absence of global changes in serum free fatty acids (FFA), triglycerides (TG), glucose, body weight or lean body mass (Figure 5A-F).

**Figure 5.**
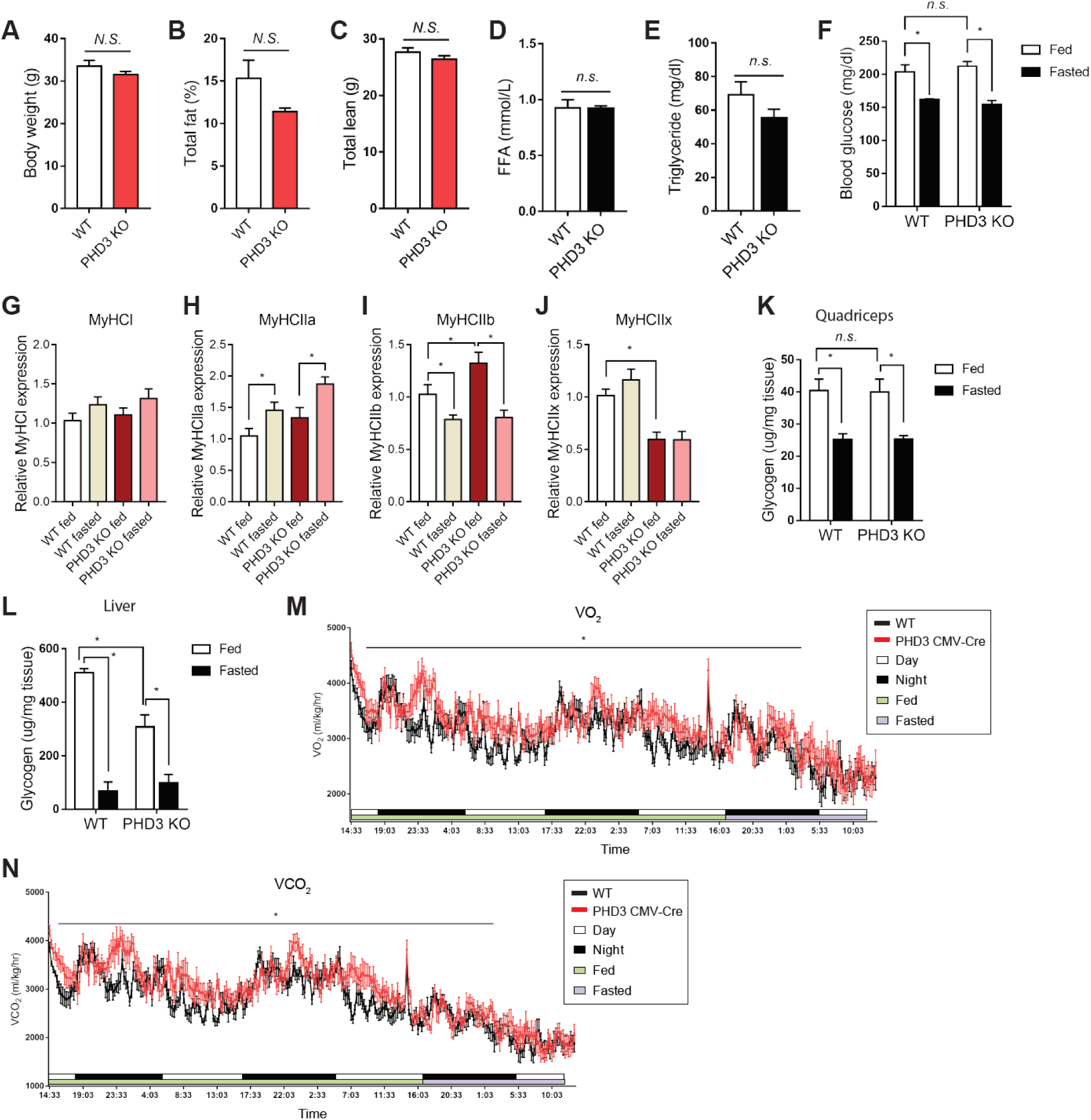
Characterization of PHD3 KO mice. Metabolic parameters in 20 weeks of WT or PHD3 KO mice using DEXA imaging analysis (n = 9 mice per group). Body weight (A), total fat (B), total lean mass (C) in WT or PHD3 KO mice (n=9). Blood FFA (D) and triglyceride (E) levels using blood chemistry (n=4). (F) Blood glucose levels in WT or PHD3 KO mice under fed or fasted condition (n=9 per group). The mRNA levels of MyHCI (G), MyHCIIa (H), MyHCIIb (I) and MyHClIx (J) in WT or PHD3 KO mouse quadriceps under fed or fasting conditions (n = 4). Glycogen levels were measured and normalized by tissue weight of WT or PHD3 KO mice quadriceps (K) or liver (L) under fed or fasted condition (n=4). (M) Mean whole-body maximum oxygen consumption rate (VO2) in 20 weeks of WT or PHD3 KO mice using Comprehensive Lab Animal Monitoring System (CLAMS) (n=9). Animals were fed ad libitum for 48 hours and fasted for 16 hours. Data represented means ± SEM. *P < 0.05. (N) Mean whole-body CO2 analysis (n=9).

### Loss of PHD3 increases exercise capacity

Since PHD3 loss activates FAO *in vitro* and *in vivo*, and ACC2 and PHD3 are highly expressed in oxidative tissues (Figure 2A), we sought to identify physiological conditions where PHD3 function may be important. In an unbiased analysis using a publicly available human skeletal muscle dataset with 242 samples (GSE3307) (Bakay et al., 2006; Dadgar et al., 2014), PHD3 expression negatively correlated with mitochondrial and FAO associated genes (Figure S5A). However, PHD3 expression did not appear to significantly change in a datasets from human skeletal muscle biopsies either obtained from a post-exercise cohort in humans (GSE111551) (Figure S6B) or after a short endurance exercise challenge in mice (Figure S6C).

Next, we investigated parameters of skeletal muscle function and physiology. We investigated whether PHD3 KO mice displayed changes in quadriceps muscle fiber type by measuring the gene expression of Myosin heavy chain 1 (MyHCI) and Myosin heavy chain 2 (MYH2), which are type I and type II fiber markers, respectively (Figure 5G-J) (Bloemberg and Quadrilatero, 2012). Compared with control animals, PHD3 KO mice displayed similar expression of MyHCI, MyHCIIa, MyHCIIb and MyHCIIx (Figure 5G-J). We also examined the gene expression of muscle differentiation markers MyoD which were increased, and myoglobin which were not changed in PHD3 KO quadriceps compared to WT tissue (Figure S5D-F). Moreover, there was no change in MYH2 and ACC2 protein level in PHD3 and WT quadriceps using immunofluorescence staining (Figure S5).

Distinct from reported mitochondrial biogenesis associated with AMPK activation (Narkar et al., 2008), PHD3 KO mouse quadriceps showed a lack of global induction of TFB1M, TFB2M, PGC1α, cytochrome C, PDK4, and PPARα (Figure S5I). As glycogen serves as a major source of stored energy, we also measured quadriceps and liver glycogen levels (Figure 5K-L). The level of glycogen in quadriceps was unchanged in PHD3 KO mice compared with WT animals. In the liver, glycogen levels were reduced in fed PHD3 KO mice, and became similarly low upon fasting. Finally, we examined whole body energy homeostasis in PHD3 KO animals by measuring the oxygen consumption (VO_2_), CO_2_ release (VCO_2_) and respiratory exchange ratio (RER) of WT and PHD3 KO mice in fed versus fasted states (Figure 5M-N, S5J). VO_2_ was increased in PHD3 KO mice during the comprehensive lab animal monitoring system (CLAMS) compared to WT (Figure 5M), and VCO_2_ was increased in PHD3 KO mice. As a result, RER was not significantly changed in PHD3 KO animals compared to WT (Figure 5N and S5J).

During exercise, muscle cells oxidize fatty acids, in addition to carbohydrates, to support energy demand (Spriet, 2014; Mul et al., 2015). Previous studies have shown that AMPK increases body energy expenditure though phosphorylation of ACC2 (O’Neill et al., 2014; Galic et al., 2018). Loss of ACC2 increases exercise capacity (Choi et al., 2007; Cho et al., 2010; Abu-Elheiga et al., 2003), and AMPK activators, such as AICAR, have been shown to increase exercise tolerance by up to 44%, by inducing mitochondrial gene expression and biogenesis (Narkar et al., 2008). Thus, we reasoned that PHD3 loss may be important physiologically during the challenge of exercise tolerance. Based on our biochemical, metabolic, and *in vivo* data supporting a reciprocal interaction between AMPK and PHD3, we hypothesized that PHD3 KO mice would possess increased energy expenditure and exercise capacity, and may phenocopy these aspects of AMPK activation *in vivo*. To test this idea, we analyzed exercise endurance in control and PHD3 KO mice using a metabolic treadmill with a gradually increasing incline (Figure 6A, Figure S6A). Remarkably, during the exercise endurance challenge PHD3 KO mice displayed a significantly increased VO_2_, particularly during the high incline period (Figure 6B); thus, loss of PHD3 function enabled animals to maintain elevated VO_2_ during rigorous exercise. Consistent with elevated metabolic fitness endurance, PHD3 KO animals generated more VCO_2_ (Figure S6C). PHD3 KO mice ran for 40% longer time and >50% further distance compared with WT animals (Figure 6C-E). The maximal oxygen consumption rate (VO_2max_) was unchanged between WT and PHD3 KO animals (Figure S6B). However, WT mice reached VO_2max_ sooner than PHD3 KO animals, consistent with the finding that PHD3 KO mice had more overall exercise endurance (Fig 6E).

**Figure 6.**
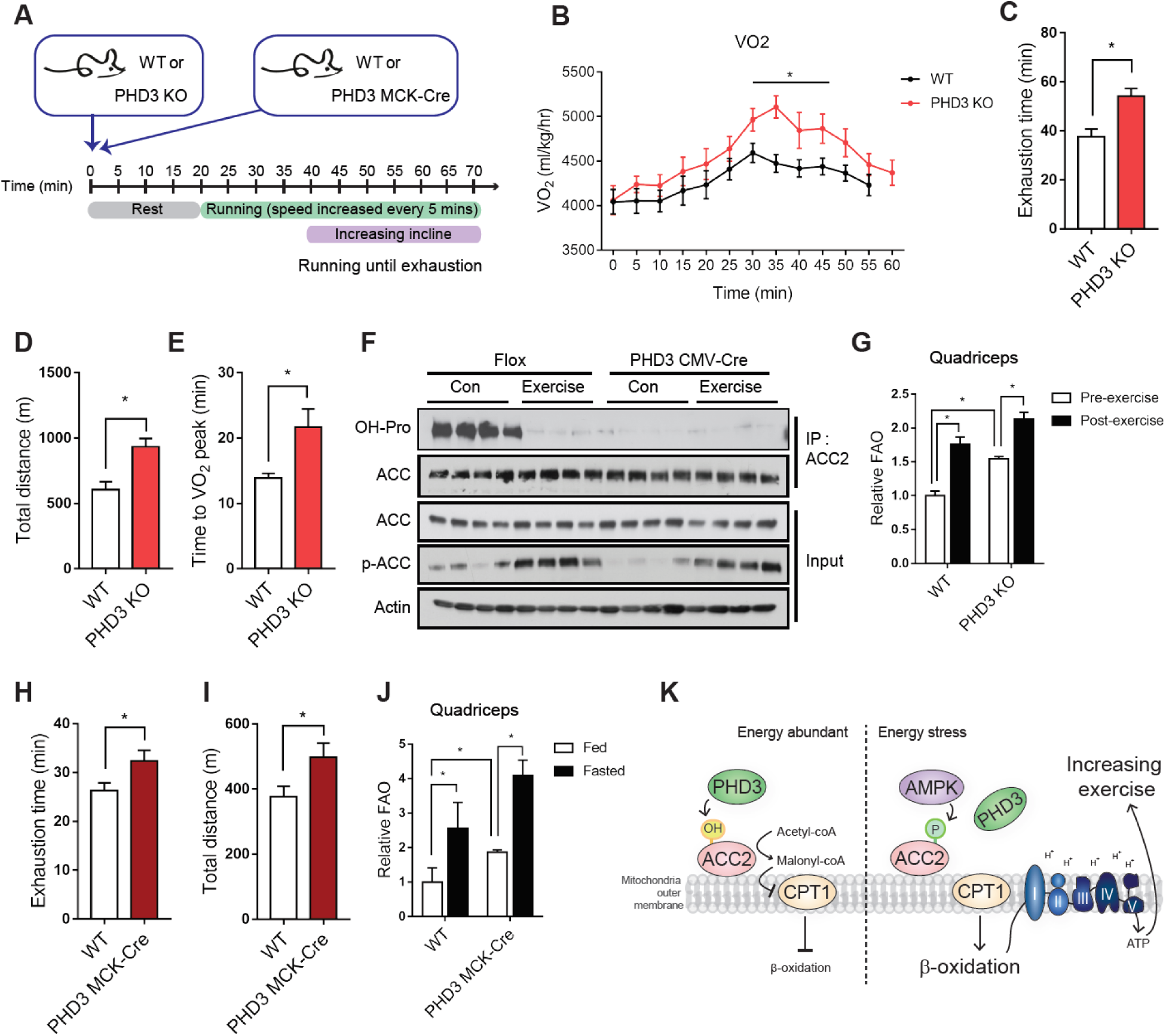
Loss of PHD3 increases exercise capacity. (A) Flowchart of the endurance exercise experiments. (B) PHD3 KO mice demonstrated increased exercise tolerance compared to WT control mice after repetitive treadmill running (n = 8 mice per group). Mean whole-body maximum oxygen consumption rate of both genotypes was calculated from the individual performances during treadmill running. The individual exhaustion time (C), total running distance (D) and time to maximum oxygen consumption rate (E). Data represent means ± SEM. *P < 0.05. (F) Immunoprecipitation and western blotting analysis of ACC2 hydroxylation and phosphorylation in WT or PHD3 KO mouse quadriceps under control or exercise conditions (n = 4). (G) Palmitate oxidation in WT or PHD3 KO mice quadriceps under pre- or post-exercise conditions using 14C-labeled palmitate and CO2 capture FAO analysis (n=3). The individual exhaustion time (H), total distance for running (I) during treadmill running with WTFL and PHDS muscle specific knock-out mice (PHD3 MCK-Cre). (J) Palmitate oxidation in control or PHD3 MCK-Cre mice quadriceps under fed or fasted condition using 14C-labeled palmitate and CO2 capture FAO analysis (n=4). (K) Schematic that loss of PHD3 induces exercise capacity. PHD3 hydroxylates ACC2 to represses FAO and AMPK induces FAO by inhibiting PHD3 binding into ACC2.

Mechanistically, we tested whether ACC2 hydroxylation changed in exercised skeletal muscle. As expected, loss of PHD3 decreased ACC2 hydroxylation in quadriceps muscle (Figure 2E, Figure 6F). Strikingly, while ACC2 protein levels were unchanged, ACC2 hydroxylation was nearly undetectable in skeletal muscle isolated from mice immediately after endurance exercise. This hydroxylation was dependent upon PHD3, as PHD3 KO mice demonstrated little to no ACC2 hydroxylation in either condition. As expected, the phosphorylation of ACC2 was increased by exercise, and was equally induced with similar kinetics in skeletal muscle from PHD3 KO animals (Figure 6F). To examine PHD3-dependent FAO in exercise, we measured the rate of ^14^C-palmitate oxidation on quadriceps slices isolated from WT and PHD3 KO mice after endurance exercise. Parallel with hydroxylation changes, *ex vivo* FAO was increased in WT mice quadriceps post-exercise compared to pre-exercise (Figure 6G). Furthermore, the quadriceps of PHD3 KO mice displayed higher FAO compared to WT (Figure 6G). Interestingly, PHD3 KO mouse quadriceps showed increasing gene expression of VEGF, TFB1M, TFB2M, PGC1α, PDK4, and PPARα after endurance exercise compared to WT animals (Figure S6E).

To test whether PHD3 loss of function in skeletal muscle was sufficient to increase exercise capacity, we generated MCK-Cre driven PHD3 knockout mice (PHD3 MCK-Cre), to delete PHD3 specifically in muscles. Indeed, PHD3 levels were decreased in quadriceps of PHD3 MCK-Cre mice compared to WT (Figure S6F) muscle, but were unchanged in the liver. When subjected to an endurance exercise challenge, PHD3 MCK-Cre mice remained on the treadmill for 23% longer time and ran 32% further distance compared with WT animals (Figure 6H-I). Consistent with our findings in cells and in tissues from PHD3 whole body KO animals, ACC2 hydroxylation was nearly undetectable in PHD3 MCK-Cre quadriceps muscle, but readily detectable in control WT tissue (Figure S6G). Next, we measured FAO from *ex vivo* slices of WT or PHD3 MCK-Cre quadriceps (Figure 6J). In both WT and PHD3 MCK-Cre quadriceps, FAO was increased after a 16-hour fast compared to fed conditions. However, PHD3 MCK-Cre quadriceps displayed elevated FAO compared with WT quadriceps (Figure 6J). In sum, these results demonstrate that PHD3 loss specifically in skeletal muscle is sufficient for increased exercise capacity.

## DISCUSSION

Here, we elucidate a clear role for PHD3 in the control of fatty acid oxidation during energetic stress. AMPK and PHD3 both appear to ACC2 in reciprocal times. During energetic stress, AMPK phosphorylates ACC2, and we observe that this phosphorylation interferes with the interaction between ACC2 and PHD3, consequently inhibiting ACC2 hydroxylation (Figure 6K). Importantly, hydroxylation of ACC2 does not affect AMPK binding to or phosphorylation of ACC2. Thus, PHD3 appears to counterbalance AMPK signaling to regulate fatty acid metabolism in response to changing nutrient availability. During physiological energy challenges, such as fasting or exercise endurance, PHD3 KO mice display greater oxidative metabolism and increased exercise capacity.

How do PTMs control ACC2 activity? First, oligomerization is a key regulator of ACC2 function; the monomeric form is inactive while the dimerized and multi-oligomerized form is active. Phosphorylation of S222 shifts ACC2 into the inactive monomeric form, demonstrating that ACC2 PTMs can affect oligomerization state. Interestingly, recent studies have shown that oligomerization of ACC2 is mediated by the BC domain, which contains the P450 residue that is hydroxylated by PHD3 (Wei and Tong, 2014; German et al., 2016). S222 and P450 are not adjacent in the amino acid sequence; yet, when the human ACC2 biotin carboxylase domain (PDB: 3JRW) (Cho et al., 2010) was modeled and superimposed with the cryo-electron microscopy ACC structure from yeast (PDB: 5CSL) (Wei and Tong, 2015), the distance between these residues may be only ∼ 50 Å (Figure 3A). Further biochemical studies are required to understand whether there is any significance to the potential proximity of S222 and P450. Importantly, our studies do not rule out a scenario whereby AMPK signaling, as well as other energy sensitive pathways alter PHD3 activity via indirect effects on metabolism. It will be interesting for future studies to determine the precise mechanism by which PHD3 influences ACC2 activity. Likewise, it has been reported that HIF signaling represses the expression of genes important in fatty acid catabolism (Liu et al., 2014; Du et al., 2017; Huang et al., 2014). Thus, multiple mechanisms may work in concert.

In this study, we determine that PHD3 loss results in increased exercise capacity. ACC2 phosphorylation by AMPK is increased during energy stresses, such as fasting, exercise, or high fat diet challenge (Cho et al., 2010; O’Neill et al., 2014; Gwinn et al., 2008; Vasquez et al., 2008; Mihaylova and Shaw, 2011). How does PHD3 loss increase exercise endurance? We demonstrate that whole body PHD3 loss enables mice to maintain high VO_2_ during an exercise challenge and run >50% further on an incline (Figure 6B and 6D). As muscle and liver glycogen levels are not elevated, increased glycogen stores could not account for the increase in exercise capacity in PHD3 KO animals (Figure 5F, 5K-L). It will be interesting for future studies to identify direct and indirect mechanisms by which PHD3 controls energy performance, such as whether this involves other AMPK mechanisms or control of HIF signaling. Interestingly, our data shows that ACC2 hydroxylation in heart tissue is profoundly responsive to fed/fasting status (Figure 2C). Therefore, these changes in exercise performance also might be related to cardiac function or involve other tissues. Importantly, increased exercise capacity also was observed in mice with muscle-specific deletion of PHD3, demonstrating that PHD3 loss in skeletal muscle is sufficient for the observed increased exercise phenotype.

Our results reveal that PHD3 and AMPK reciprocally regulate fat metabolism and exercise capacity. During energy stress, AMPK-mediated phosphorylation is sufficient to inhibit PHD3 interaction with and hydroxylation of ACC2. Upon energy replete conditions, AMPK is inactive, ACC is not phosphorylated, and PHD3 is able to hydroxylate ACC2 (Figure 6K). Collectively, these data suggest that PHD3 loss or inhibition could provide a strategy for treatment of a wide range of metabolic diseases, and that modulation of PHD3 may improve energy homeostasis and skeletal muscle performance during exercise challenge.

## MATERIALS AND METHODS

Detailed methods are provided in this paper and include the following:

- KEY RESOURCES TABLE
- CONTACT FOR REAGENT AND RESOURCE SHARING
- EXPERIMENTAL MODEL AND SUBJECT DETAILS

- Cell Lines
- Mice
- METHOD DETAILS

Preparation of plasmids, siRNAs and transfection
Quantitative RT-PCR analysis
Immunoprecipitation, western blotting and antibodies
*In vitro* protein binding assay
Immunofluorescence
Molecular modeling
Fatty acid oxidation measurements
MT-DNA quantification
Mitochondrial mass quantification
Respiration
Metabolite profiling and mass spectrometry
GeLC-MS/MS
MALDI MSI
Exercise performance studies
Histological procedures
Bioinformatics analysis
- QUANTIFICATION AND STATISTICAL ANALYSIS

### KEY RESOURCES TABLE

| REAGENT or RESOURCE | SOURCE | IDENTIFIER |
| --- | --- | --- |
| Antibodies |  |  |
| Anti-ACC | Cell Signaling Technologies | Cat# 3676S |
| Anti-ACC2 | Cell Signaling Technologies | Cat# 8578S |
| Anti-phospho-ACC | Cell Signaling Technologies | Cat# 3661S |
| Anti-AMPKα | Cell Signaling Technologies | Cat# 2532S |
| Anti-AMPKα (Thr172) | Cell Signaling Technologies | Cat# 2535S |
| Anti-PHD3 | Novus Biologicals | Cat# NB100-139 |
| Anti-PHD3 | Santa Cruz Biotechnology | Cat# sc-46030 |
| Anti-hydroxyproline | Abcam | Cat# ab37067 |
| Anti-HA | Cell Signaling Technologies | Cat# 3724S |
| Anti-HA | Cell Signaling Technologies | Cat# 2367S |
| Anti-Flag | Sigma-Aldrich | Cat# F2555 |
| Anti-GST | Santa Cruz Biotechnology | Cat# sc-138 |
| Anti-His | MLB | Cat# PM032 |
| Anti-Actin | Sigma-Aldrich | Cat# A2066 |
| Anti-α-Tubulin | Santa Cruz Biotechnology | Cat# sc-8035 |
| Anti-MYH2 | Developmental Studies<br>Hybridoma Bank | Cat# 2F7 |
| EZview™ Red Protein G Affinity Gel | Sigma-Aldrich | Cat# E3403 |
| EZview™ ANTI-FLAG® M2 Affinity Gel | Sigma-Aldrich | Cat# F2426 |
| Glutathione Sepharose 4B | GE Healthcare Life Sciences | Cat# 17075601 |
| Pierce™ HisPur™ Ni-NTA Resin | Thermo Scientific | Cat# 88221 |
| Goat anti-Rabbit IgG (H+L) Cross-Adsorbed Secondary Antibody, Alexa Fluor 488 | Invitrogen | Cat# A-11008 |
| Rabbit anti-Goat IgG (H+L) Cross-Adsorbed Secondary Antibody, Alexa Fluor 488 | Invitrogen | Cat# A-11078 |
| Goat anti-Mouse IgG (H+L) Cross-Adsorbed Secondary Antibody, Alexa Fluor 594 | Invitrogen | Cat# A-11032 |
| Goat anti-Rabbit IgG (H+L) Cross-Adsorbed Secondary Antibody, Alexa Fluor 594 | Invitrogen | Cat# A-11012 |
| Biological Samples |  |  |
| Skeletal muscle (frozen) taken from experimental mouse strains listed in the Experimental Model section | See Experimental Models | N/A |
| Hearts (frozen) taken from experimental mouse strains listed in the Experimental Model section | See Experimental Models | N/A |
| Livers (frozen) taken from experimental mouse strains listed in the Experimental Model section | See Experimental Models | N/A |
| Chemicals, Peptides, and Recombinant Proteins |  |  |
| Mitotracker Green | Invitrogen | Cat# M22425 |
| DAPI (4',6-Diamidino-2-Phenylindole, Dihydrochloride) | Sigma-Aldrich | Cat# D9542 |
| Dimethyl sulfoxide | Sigma-Aldrich | Cat# D2650 |
| Dimethyloxaloylglycine | Santa Cruz Biotechnology | Cat# sc-200755 |
| AICAR | Cell Signaling Technologies | Cat# 9944S |
| MG132 | R&D Systems INC | Cat# 1748/5 |
| LY294002 | Cell Signaling Technologies | Cat# 9901S |
| L-Carnitine | Sigma-Aldrich | Cat# C0158 |
| Palmitic acid | Sigma-Aldrich | Cat# P0500 |
| 1 μCi [9,10(n)-<sup>3</sup>H]palmitic acid | Perkin Elmer | Cat#<br>NET043001MC |
| Palmitate-BSA FAO Substrate | Agilent Technologies | Cat# 102720-100 |
| Hydrogen peroxide solution 30% | Sigma-Aldrich | Cat# H1009 |
| Western Lightning ECL Pro | Perkin Elmer | Cat#<br>NEL120001EA |
| ECL Plus Western Blotting Substrate | Thermo Scientific | Cat# 32132 |
| Commercial Assays |  |  |
| Liquid DAB+ Substrate Chromogen System | DAKO | Cat# K346811-2 |
| VECTASTAIN Elite ABC Kit | Vector Laboratories | Cat# PK-6100 |
| Experimental Models: Cell Lines |  |  |
| HEK293T | ATCC | Cat# CRL-11268 |
| Immortalized MEF | ATCC | Cat# CRL-2907 |
| C2C12 | ATCC | Cat# CRL-1772 |
| AMPKα KO and WT MEFs | From the laboratory of Dr. Reuben Shaw | N/A |
| Experimental Models: Organisms/Strains |  |  |
| Mouse: AMPK<sup>FL</sup>;Ubc-Cre<sup>ER</sup> | From the laboratory of Dr. Reuben Shaw | N/A |
| Mouse: PHD3 MCK-Cre, PHD3-CMV-Cre and PHD3-flox | This paper | N/A |
| Mouse: C57BL/6 | Jackson Laboratory | Cat# 000664 |
| Oligonucleotides |  |  |
| qPCR primers: see table S1 | This paper | N/A |
| MT-DNA copy number primers: see table S1 | This paper | N/A |
| shRNA- hairpin primers: see table S1 | This paper | N/A |
| Primers for cloning: see table S1 | This paper | N/A |
| Recombinant DNA |  |  |
| Plasmid: pcDNA3.1-HA-PHD3 | From the laboratory of Dr. William G.Kaelin Jr. lab | N/A |
| Plasmid: HA-pBabe PHD3 | From the laboratory of Dr. William G.Kaelin Jr. lab | N/A |
| Plasmid: pcDNA3.1-ACC2 WT-Flag, pcDNA3.1-ACC2 P450A-Flag, pcDNA3.1-ACC2 S222A-Flag, pcDNA3.1-ACC2 P450A/S222A-Flag, pcDNA3.1-ACC2 S222E-Flag, pcDNA3.1-ACC2 NT-Flag | This paper | N/A |
| Plasmid: pcDNA3.1-PHD3-HA | This paper | N/A |
| Plasmid: pHAGE-CMV-Flag-HA-puromycin | From the laboratory of Dr. William G.Kaelin Jr. lab | N/A |
| Plasmid: pHAGE-CMV-ACC2 WT-Flag-HA, pHAGE-CMV-ACC2 P450A-Flag-HA, pHAGE-CMV-ACC2 S222A-Flag-HA, pHAGE-CMV-ACC2 P450A/S222A-Flag-HA | This paper | N/A |
| Plasmid: pET28-PHD3-His-SUMO | This paper | N/A |
| Plasmid: pGEX-6P-1-ACC2 BC, pGEX-6P-1-ACC2 BCCP, pGEX-6P-1-ACC2 CT | This paper | N/A |
| Plasmid: pLKO.1-puro, pLKO.1-shPHD3 #1, pLKO.1-shPHD3 #2 | This paper | N/A |
| Plasmid: pLKO.1-puro, pLKO.1-shACC2 #1, pLKO.1-shACC2 #2 | This paper | N/A |
| Software and Algorithms |  |  |
| ImageJ | NIH | N/A |
| GraphPad Prism 7.0 | GraphPad Software | N/A |

### CONTACT FOR REAGENT AND RESOURCE SHARING

Further information and requests for resources and reagents should be directed to and will be fulfilled by the Lead Contact, Marcia Haigis.

### EXPERIMENTAL MODEL AND SUBJECT DETAILS

#### Cell Lines

HEK293T (human embryonic kidney), C2C12 (mouse mesenchymal precursor) and immortalized MEF cell-lines were obtained from American Type Culture Collection (ATCC). Cells were cultured at 5% CO_2_ in 4.5 g/L (25 mM) glucose Dulbecco’s modified Eagle’s medium, supplemented with 10% fetal bovine serum and penicillin/streptomycin. C2C12 differentiation toward myoblast was induced by 2% horse serum (Gibco) and Insulin-Transferrin-Selenium Liquid Media Supplement (Sigma) for two days. AMPK KO MEFs and WT controls were obtained from Reuben Shaw at the Salk Institute, as described (Gwinn et al., 2008).

#### Mice

AMPK^FL^;Ubc-Cre^ER^ mice were from the laboratory of Reuben J. Shaw (The Salk Institute for Biological Studies). AMPK^FL^;Ubc-Cre^ER^ mice were generated by crossing mice with floxed AMPKα1 and AMPKα2 alleles (provided by Benoit Viollet) to Ubc-Cre^ER^ mice (Ruzankina et al., 2007). Mice were backcrossed into a FVB/n background. Experiments were started when mice were 8 weeks of age. Only male mice were used. To induce AMPK deletion, AMPK^FL^;Ubc-Cre^ER^ mice were injected (intraperitoneal injection) with tamoxifen (Sigma) for 5 consecutive days (4 mg tamoxifen per day, suspended in 200 μl corn oil + 2% ethanol). Muscle tissue was collected 3 weeks after tamoxifen treatment and snap frozen in liquid nitrogen. For PHD3-CMV-Cre mice, Egln2*^tm2Fong^* Egln1*^tm2Fong^* Egln3*^tm2Fong^*/J mice (PHD1^FL^/PHD2^FL^/PHD3^FL^triple floxed strain # 028097) (Takeda et al., 2006) were obtained from The Jackson Laboratory and housed in the New Research Building Animal Facility at Harvard Medical School. PHD3^FL^ mouse were obtained by breeding between Egln2*^tm2Fong^* Egln1*^tm2Fong^* Egln3*^tm2Fong^*/J mice and C57BL/6J. PHD3^FL^ mouse were crossed with B6.C-Tg(CMV-cre)1Cgn/J mice (CMV-Cre mice # 006054) to generate PHD3-CMV-Cre knock-out mice. To generate PHD3 MCK-Cre knock-out mice, PHD3^FL^ mouse were crossed with B6.FVB(129S4)-Tg(Ckmm-cre)5Khn/J mice (MCK-Cre mice # 006475). Genotyping was performed by PCR using genomic DNAs obtained from the tails. All animal studies were performed in accordance with protocols approved by the Institutional Animal Care and Use Committee, the Standing Committee on Animals at Harvard.

### EXPERIMENTAL METHOD DETAILS

#### Preparation of plasmids, siRNAs and transfection

pcDNA3.1-HA-PHD3 or HA-pBabe PHD3 were previously described (Lee et al., 2005). ACC2 cDNA in pENTR223 vector was obtained from the Dana Farber/ Harvard Cancer Center Resource Core. ACC2 cDNA was cloned into pHAGE-CMV-Flag-HA-puromycin vector (obtained from Wade Harper’s lab at Harvard Medical School) using Gateway LR Clonase II Enzyme Mix according to manufacturer’s instructions. For cloning, 10 μl reactions containing 150 ng ACC2 pENTR223, 150 ng pDEST vector and 2 μl Clonase in TE buffer (pH 8.0) were incubated at 25°C for 2 hours. Site-specific mutations of ACC2 and stop codon deletion constructs were performed using PCR-based mutagenesis (QuikChange II XL Site-Directed Mutagenesis Kit, Agilent). The plasmids for C-terminus GST-tagged or Flag-tagged ACC2 fragments were re-cloned from the human ACC2 plasmid using PCR and Gibson assembly. For transient transfection with plasmids, cells at ∼40% density were transfected with plasmids using Fugene HD (Roche). The transfected cells were cultured for 48 hours, and then selected with puromycin. The shRNAs containing a hairpin loop were synthesized and inserted into pLKO-puromycin vector for lenti-viral infection. The viral vectors were co-transfected into HEK293T cells with pRSV-Rev, pMD2-VSVG, and pMDLg/pRRE plasmids to prepare viral particles. On the 3rd day after transfection, lentiviruses were collected from the supernatant of HEK293T cells. Cells were infected with viruses (100 ul/ml) and selected using puromycin (2 ug/ml) to establish stable cell lines.

#### Quantitative RT-PCR analysis

RNA was isolated by extraction with Trizol according to manufacturer instructions (Invitrogen) or with the RNA Clean & Concentrator Kit (Zymo Research). cDNA was synthesized using iScript cDNA synthesis kit (BioRad). Quantitative real-time PCR was performed with SYBR Green Fast Mix (Quanta Biosciences) on a Roche Lightcycler 480 and analyzed by using ΔΔCt calculations. qPCR analyses in human cell lines are relative to the reference gene B2M. qPCR analyses in mouse cell line or tissue are relative to β-Actin.

#### Immunoprecipitation, western blotting and antibodies

Western blotting was performed using antibodies against ACC (Cell Signaling Technologies (CST), ACC2 isoform (CST), actin (Sigma), AMPKα (CST), HA (CST), Flag (Sigma), hydroxyproline (Abcam), phospho-ACC (CST), phospho-AMPKα (CST), PHD3 (Novus Biologicals) and tubulin (Sigma). 1% NP40 buffer containing protease and phosphatase inhibitors was used to prepare lysates, unless otherwise indicated. For ACC hydroxylation time course studies, the pan-PHD inhibitor dimethyloxalylglycine (DMOG, Frontier Scientific, 1 mM) was added to the lysis buffer to prevent further hydroxylation in the lysate. For immunoprecipitations of transiently overexpressed HA-tagged or Flag-tagged proteins, lysates were immunoprecipitated using EZview anti-HA Affinity Gel (Sigma) or anti-Flag Affinity Gel (Sigma). For endogenous immunoprecipitations, lysates were immunoprecipitated with ACC antibody (CST) or ACC2 antibody (CST) and EZview Red Protein G Affinity Gel (Sigma).

#### *In vitro* protein binding assay

Recombinant ACC2-BC, ACC2-BCCP and ACC2-CT-GST peptides were expressed in *E. coli*, purified by GSH-affinity beads, and eluted by GSH, respectively. Recombinant PHD3-HA-His proteins were purified from *E. coli* using nickel NTA affinity resin. The mixtures of peptides were incubated at 4°C for 1 hour, further incubated with the affinity beads at 4°C for 4 hours, spun down, and immunoblotted.

#### Immunofluorescence

Cells cultured on glass coverslips at 40%–60% confluence were first stained by incubation with 50 nM Mitotracker Red (Thermo Fisher Scientific) in culture media for 20 min, washed, and then stained by indirect immunofluorescence. In brief, cells were rinsed in PBS and fixed with 3.7% formaldehyde in PBS for 15 min at room temperature. Cells were permeabilized with PBS plus 0.2% Triton X-100 for 10 min and blocked for 1 hour at 4°C, with PBS containing 5% normal goat serum. Cells were incubated with primary antibodies: rabbit anti-PHD3 (1:100) in PBS overnight at 4°C. After 3 washes with PBS, 5 min at room temperature, cells were incubated for 2 hours with goat anti-rabbit IgG conjugated to the fluorescent Alexa 488 dye (1:400) in PBS. After 3 washes, nuclei were detected with 1 mg/ml DAPI. After 3 washes in PBS and 1 wash with 75% ethanol, cells were mounted in ProLong Gold Antifade reagent for viewing with a confocal microscope (Nikon TE2000 w/C1 Point Scanning Confocal).

#### Molecular modeling

Using PyMol software, the biotin-carboxylase domain of human ACC2 (PDB:3JRW) (Cho et al., 2010) was superposed with the yeast cryo-electron microscopy structure of entire ACC (PDB: 5CSL) (Wei and Tong, 2015). The mapping of phosporylation site serine 222 and hydroxylation site proline 450 site were estimated and designed by sequence homology with yeast ACC and the biotin-carboxylase domain of human ACC2 (PDB: 5CSL).

#### Fatty acid oxidation measurements

For FAO assays, cells in 12 well-plates were pre-incubated 4 hours in serum-free low glucose medium supplemented with 100 μM palmitate or hexanoate and 1 mM carnitine. Cells were then changed to 600 μl medium containing 1 μCi [9,10(n)-^3^H]palmitic acid (Perkin Elmer) and 1 mM carnitine for 2 hours. The medium was collected and eluted in columns packed with DOWEX 1X2-400 ion exchange resin (Sigma) to analyze the released ^3^H_2_O. FAO in complete medium indicates medium with serum and high glucose or low glucose were used. Basal FAO indicates cells were not pre-incubated with fatty acids prior to FAO analysis. Counts per minute (CPM) were normalized to protein content in parallel cell plates.

#### MT-DNA quantification

Mitochondrial DNA was extracted from cells and analyzed by real time quantitative PCR, as described (Errichiello et al., 2015; O’Neill et al., 2011). Briefly, 50,000 cells were pelleted and re-suspended in 50 uL of MT-DNA lysis buffer (25 mM NaOH, 0.2 mM EDTA). Lysates were heated to 95°C for 15 minutes and neutralized with 50 μL MT-DNA neutralization buffer (40 mM Tris-HCl). RT-qPCR reactions were performed on 5 uL of 1:50 diluted lysate. MT-DNA quantification was performed with mitochondrial markers MT-CO2 and D-loop. MT-DNA was normalized to nuclear DNA markers β-Actin and β-Globin.

#### Mitochondrial mass quantification

50,000 cells were adapted to their respective conditons for 24 hours prior to the experiment. Cells were incubated with 75 nM Mitotracker Green (Invitrogen) for 1 hour, washed twice with PBS and analyzed on an LSR II Flow Cytometer (BD Biosciences), gated for single cells and assessed for fluorescence intensity.

#### Respiration

Respiration was assessed using the Seahorse XFe-96 Analyzer (Seahorse Bioscience). For experiments that measure the basal levels of oxygen consumption rate, shcon or shPHD3 MEF cells were incubated with 25 mM glucose or 5 mM glucose media for 8 hours prior to the experiment. Following this incubation, media was changed to a non-buffered, serum-free Seahorse Media (Seahorse Bioscience) supplemented with 10 mM glucose, 2 mM L-glutamine and 1 mM sodium pyruvate. For basal OCR using fat, cells were incubated serum free media over night and incubated serum-free Seahorse Media (Seahorse Bioscience) supplemented with 0.5 mM glucose, 1 mM L-glutamine, 0.5 mM carnitine and XF BSA-palmitate. Values were normalized to cell number.

#### Metabolite profiling and mass spectrometry

Metabolites were isolated in 80% MeOH and analyzed on two distinct methods of hydrophilic interaction liquid chromatography coupled to mass spectrometry (HILIC-MS). In one method, electrospray ionization was tailored to negative-ion mode, and in the second method to positive-ion mode. For negative-ion mode, analytes were eluted in buffer A (20 mM ammonium acetate, 20 mM ammonium hydroxide) and buffer B (10 mM ammonium hydroxide in 75:25 acetonitrile:methanol). Samples were run on a HILIC silica (3 um, 2.1 × 150 mm) column (waters) with a binary flow rate of 0.4mL/min for 10 minutes on linear gradient (95% buffer B to 0% buffer B) followed by 2 minutes with (0% buffer B) and ending with a 2 minute linear gradient (0% buffer B to 95% buffer B) and holding (95% buffer B) for 13 minutes. For positive-ion mode, samples were dried down and reconstituted in a 20:70:10: acetonitrile:MeOH:water mixture. The buffers were: buffer A (10 mM ammonium formate, 0.1% formic acid in water) and buffer B (acetonitrile, 0.1% formic acid). Samples were run on a HILIC silica (3 um, 2.1 × 150mm) column (waters) with a binary flow rate of 0.25 mL/min for 10 minutes on linear gradient (95% buffer B to 40% buffer B) followed by 4.5 minutes with (40% buffer B) and ending with a 2 minute linear gradient (40% buffer B to 95% buffer B) and holding (95% buffer B) for 13 minutes. For both ion-modes, a Q Exactive hybrid quadrupole orbitrap mass spectrometer (Thermo Fisher Scientific) with a full-scan analysis over 70–800 m/z and high resolution (70,000) was used for mass detection. A targeted-method developed for 176 compounds (118 on positive and 58 on negative) was used to identify metabolites. A master mix of reference standards for metabolites in the targeted method were run immediately prior to each set of samples, such that their retention times were associated with peaks in the unknown samples run over that same column. Peaks were integrated in Tracefinder 3.3. Metabolite levels were normalized to cell number in parallel plates or protein concentration in the same amount of tissue powder samples.

#### GeLC-MS/MS

To probe hydroxylation on proline residue, ACC2 was transiently overexpressed in shCon or shPHD3 293T cells under high or low glucose media and immunoprecipitated with ACC2 antibody (CST). Bound material was washed and separated by SDS-PAGE. The Coomassie stained band corresponding to ACC2 was analyzed by GeLC-MS/MS (Paulo, 2016). Eluted peptides were derivatized using TMT labeling (Navarrete-Perea et al., 2018) and analyzed using the MS3-IDQ method (Berberich et al., 2018). MS3 spectra were searched against the Uniprot Human database using Sequest with proline hydroxylation set as a variable modification (+15.9949 Th shift).

#### MALDI MSI

Mouse quadriceps were isolated from 8 week old mice under either fed or fasting conditions (16 hours) and snap-frozen in liquid nitrogen. Tissue samples were stored at −80°C prior to use. Muscle tissue was cryo-sectioned at a thickness of 12 µm at −20°C, tissue sections were thaw-mounted onto ITO-coated glass slides for MALDI MSI. Adjacent to the tissue on the slide were placed 0.5 µL dried droplets of standards of: myristoyl-L-carnitine, palmitoyl-L-carnitine and stearoyl-L-carnitine (10 µM, in water, Sigma). The slides were then coated with matrix (DHB, 160 mg/mL in 70/30 methanol/0.1% trifluoroacetic acid) using a TM-Sprayer (HTX imaging). Matrix was sprayed with a flow rate of 0.09 mL/min, a velocity of 1200 mm/min, track spacing of 2 mm, a nebulizer gas temperature of 75°C, a gas pressure of 10 psi and 2 passes of matrix. MALDI MSI data were acquired using a 9.4 Tesla SolariX XR FT-ICR MS (Bruker Daltonics) externally calibrated in electrospray ionization positive ion mode using a tuning mix solution (Agilent Technologies). MALDI MS images were acquired from both tissue and acylcarnitine standard spots, with a pixel step size of 75 µm. Spectra were acquired in positive ion mode with 250 laser shots accumulated at each location. Spectra were collected in the range m/z 50-3000 with isolation of ions using the constant accumulation of selected ions (CASI) mode, with Q1 set to 500 and an isolation window of 260. Online calibration of spectra to heme (*m/z* 616.17668, Δppm tolerance 200, intensity threshold 500) was applied. Also, ATP (*m/z* 500.003023, Δppm = 2.33), myristoyl-carnitine (*m/z* 372.310835, Δppm = 0.2), palmitoyl-carnitine (*m/z* 400.342135, Δppm = 1.16), and stearoyl-carnitine (*m/z* 428.373436, Δppm = 1.34) were measured. The laser focus was set to ‘small’, the laser energy was set to 35% (arbitrary scale) with a laser frequency of 1000 Hz. MALDI MS images were displayed and analyzed using FlexImaging 5.0 and SCILS Lab (2018) (both Bruker Daltonics) software. MSI data are displayed with normalization to total ion count (TIC).

#### Exercise performance studies

Mice were acclimated to the treadmill 2 days prior to the experiments by running for 5 min/day at 5 m/min and 10 m/min followed by 15 m/min for 1 min. For exercise experiment, speed was increased 5 m/min every 5 min until reaching 20 m/min, and then incline was increased 5 incline every 5 minutes until exhaustion. Respiratory exchange ratio (RER), VO_2_, VCO_2_ and heat were monitored using the Oxymax Modular Treadmill System.

#### Histological procedures

Mouse quadriceps were isolated from 8 week old mice were in fed or fasting conditions for 16 hours. Mouse quadriceps were fixed in 10% buffered formalin, dehydrated through a series of ethanol solutions of increasing concentration and submitted to the Dana-Farber/Harvard Cancer Center Pathology Cores for embedding in paraffin, sectioning, and hematoxylin and eosin staining. Immunohistostaining was performed using anti-PHD3 antibody (Novus Biologicals). The fresh frozen quadriceps were sectioned and stained with H&E. Immunofluorescence was performed on the same samples using anti-myosin heavy chain 2 (Developmental Studies Hybridoma Bank), ACC2 (Cell Signaling Technology) and DAPI. Digital images of stained sections were taken using confocal microscope (Nikon TE2000 w/C1 Point Scanning Confocal).

#### Bioinformatics analysis

Human muscle expression datasets were obtained from GEO under the respective accession identifiers and from GTEx (PMID: 23715323). Confounding factors, including gender, age, batch, and disease, as well as hidden factors that could cause gene expression variability, were estimated and removed using probabilistic estimation of expression residuals (PEER) (PMID: 22343431). Expression residuals obtained from PEER were used for further analysis. To identify the enriched gene sets correlated with PHD3 expression, we performed gene set enrichment analysis (GSEA) (PMID: 16199517) using the fgsea package (https://www.biorxiv.org/content/early/2016/06/20/060012). Specifically, genes were ranked based on the Pearson correlation coefficient against PHD3 expression, and enrichment analysis was performed to determine the enriched gene sets co-expressed with PHD3.

#### Quantification and statistical analysis

Details regarding the specific statistical tests, definition of center, and number of replicates (n), can be found for each experiment in the figure legends. All data of animal study are shown as mean ± SEM, while data from in vitro studies are shown as mean ± SD. When comparing two groups, statistical analysis was performed using a two-tailed Student’s t tests and statistical significances were considered when P values were less than 0.05. GraphPad, Prism, and Excel were used for all quantifications and statistical analyses.

## ACKNOWLEDGEMENTS

We thank all members of the Haigis lab for thoughtful discussion on this manuscript. H.Y. is funded by the American Diabetes Association Fellowship (1-17-PDF-109) and M.C.H is funded by NIH grant R01CA213062, the Ludwig Center at Harvard University, and Glenn Foundation for Medical Research. J.A. is funded by the Ecole Polytechnique Fédérale de Lausanne and the Fondation Suisse de Recherche sur les Maladies Musculaires. A.C., A.D., L.J.G is funded by the Joslin Core grant (NIH P30DK036836). E.C.R. is in receipt of an NIH R25 (R25 CA-89017) Fellowship in partnership with the Ferenc Jolesz National Center for Image Guided Therapy at BWH (P41 EB015898). H.Y. and M.C.H are co-inventors on the patent case number-HU 7395 submitted by Harvard Medical School that Dynamic regulation of fat metabolism by Acetyl CoA Carboxylase. Microscopy experiments were performed at the Nikon Imaging Center at Harvard Medical School. Mouse exercise experiments were performed at Joslin Diabetes Center. pHAGE-CMV-Flag-HA-puromycin vector was obtained from Wade Harper’s lab at Harvard Medical School. cDNA and entry vectors were obtained from the Dana Farber/ Harvard Cancer Center Resource Core.

## AUTHOR CONTRIBUTIONS

H.Y. and M.C.H. designed experiments, wrote the paper, and analyzed data. H.Y, J.B.S., P.J.A., and S.P.G performed and analyzed mass spectrometry experiments. H.Y., E.Z., and S.J.W. conducted *in vivo* mouse experiments. E.C.R performed mass spectrometry imaging and data analysis, N.Y.R.A. contributed analytical tools. H.Y., A.D., A.C., and L.J.G performed mouse exercise experiment in Joslin Diabetes Center. H.Y., E.Z., S.J.W, Y.W., and N.J.G performed experiments on gene expression, protein levels and post-translational modification. D.G., and R.J.S provided tissue samples for AMPK KO mice. H.Y. H.L., and J.A. performed bioinformatics analysis. H.Y. performed structure modeling.

## SUPPLEMENT INFORMATION

**Figure S1.**
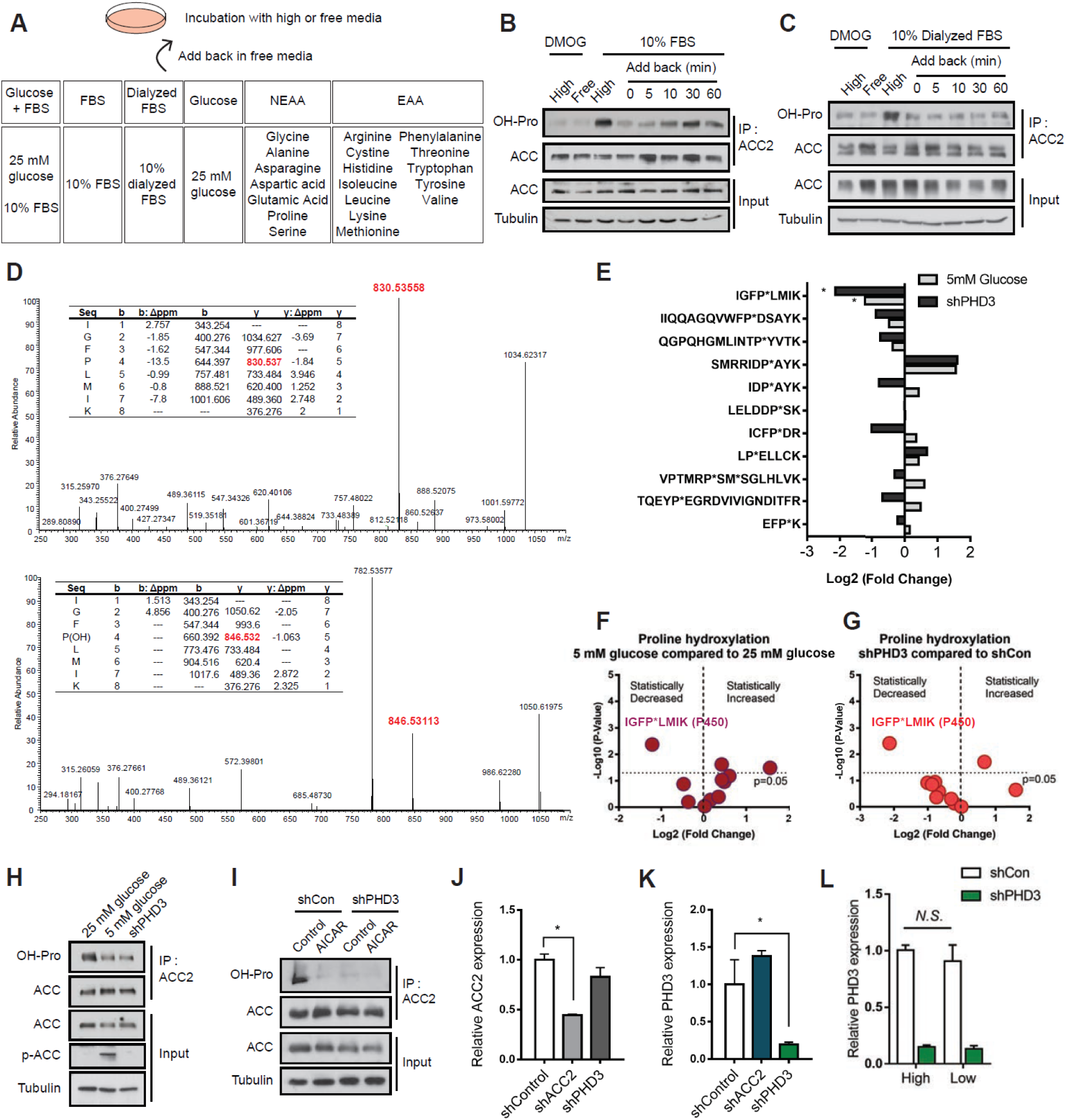
The hydroxylation of proline 450 residue is most changed by nutrient-depleted conditions as well as PHD3 depletion compared to high nutrient conditions. (A) Schematic depicting of the different fuels to which cells in culture are exposed. After fuel addition, cells were used for ACC2 hydroxylation assay. Hydroxylation was assessed after 12 h incubation in high or nutrient-free media, and then 0 to 60 min after 10% FBS (B), and 10% dialyzed FBS (C) re-addition. Hydroxylation was detected by western blotting. (D) Representative mass spectra identifying the hydroxylated and non-hydroxylated versions of residue P450 in pulled down ACC2 protein. ‘‘b’’ fragments contain the N-terminal amino acid and are labeled from the N to C terminus. ‘‘y’’ fragments contain the C-terminal amino acid and are labeled from the C to N terminus. (E) Relative abundance of hydroxylation of proline residues on ACC2 in 5 mM glucose media incubated (dark grey) or depleted of PHD3 (light grey) compared to control cells. For all comparisons two-tailed t test was used. P < 0.05. Volcano plots demonstrating that 5 mM glucose media (F) or PHD3-depleted MEFs (G) incubated MEFs have statistically significant proline hydroxylation changes compared to control cells. The proline 450 residue, the sequence of which is IGFP(OH)LMIK, was most significantly changed. (H) Immunoprecipitation and western blotting analysis of ACC2 hydroxylation and phosphorylation in indicated condition. shControl or shPHD3 293T cells were transfected with ACC2, and incubated with 25 mM glucose media or 5 mM glucose media for 4 h. ACC2 proteins were pulled down using antibody against ACC2 and immunoblotted using the indicated antibodies. (I) Immunoblots of ACC2 hydroxylation from MEFs expressing either control shRNA or PHD3 shRNA with or without AICAR. (J) Relative ACC2 expression using mouse β-actin as a reference gene in MEFs expressing shRNA against ACC2, PHD3 or shRNA control (n = 4). (K) Relative PHD3 expression using mouse β-actin as a reference gene in MEFs expressing shRNA against ACC2, PHD3 or shRNA control (n = 4). (L) PHD3 expression using mouse b-actin as a reference gene in MEFs under high or low nutrient conditions.

**Figure S2.**
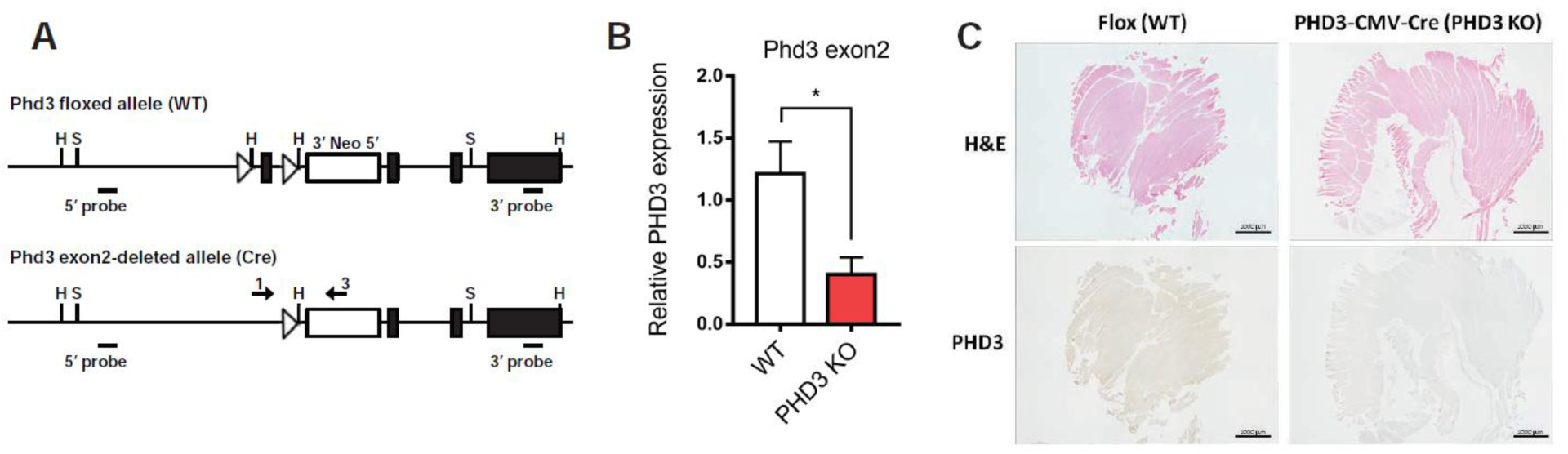
Estabishment of PHD3 KO mice. (A) Gene targeting strategy for Phd3. In targeted Phd3 locus, exon 2 is flanked by two loxP sites. Exon 2 is deleted by Cre-mediated recombination. S, Sca I; H, Hind III; Neo, neomycin cassette; hatched square, Frt site (flanking the Neo cassette); open triangle, loxP site; arrows 1, 2 and 3. (B) qPCR analyses to confirm the lack of WT mRNA transcripts in PHD3 KO mouse RNA samples. (C) H&E and immunohistochemistry staining of PHD3 in WT or PHD3 KO mice muscles. Scale is 100 μm.

**Figure S3.**
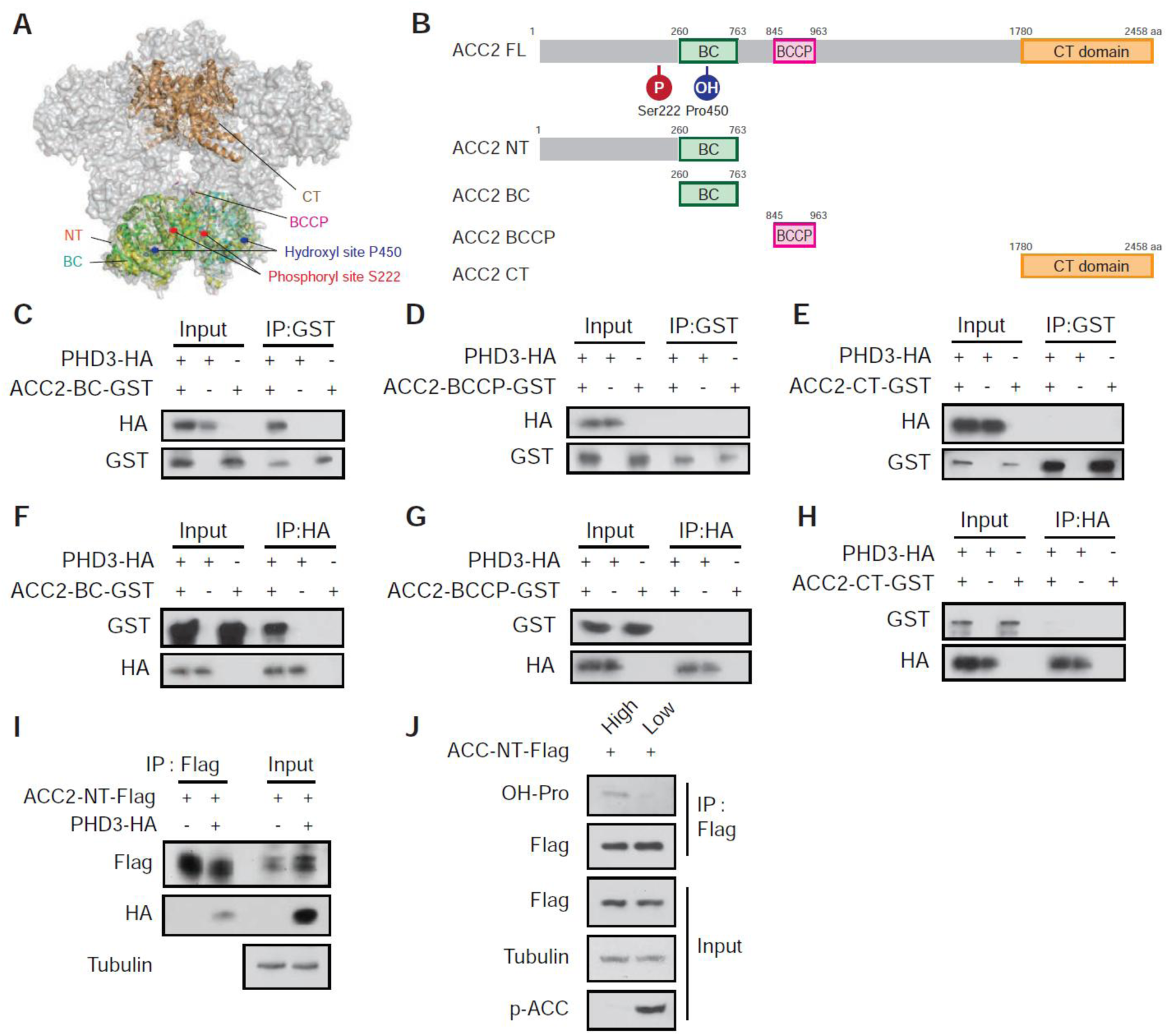
PHD3 binds the BC domain of ACC2. (A) P450 (hydroxylation site, blue) is located in the ACC2 BC domain (green and cyan). S222 (phosphorylation site, red) is located inside of the ACC holoenzyme dimer structure. This model was generated by superposition of human ACC2 biotin carboxylase domain (PDB: 3JRW) with the entire yeast ACC (PDB: 5CSL). (B) Schematic figure of the domains of ACC2 protein. Posttranslational modifications by AMPK (phosphorylation, red) and PHD3 (hydroxylation, blue) are highlighted. BC, biotin carboxylase domain; BCCP, biotin carboxyl carrier protein domain; CT, carboxyltransferase domain; N-terminus domain contains ATP grasp domain. (C) Immunoprecipitation and western blotting analysis of PHD3 interaction with the BC domain of ACC2. Proteins were expressed in E. coli. ACC2 BC domain with GST-tag and PHD3-HA was pulled down using HA-beads for an in vitro binding assay, and the precipitates were immunoblotted. (D) In vitro binding assay of PHD3 interaction with the BCCP domain using HA-beads. (E) In vitro binding assay of PHD3 interaction with the CT domain using HA-beads. (F) Immunoprecipitation and western blotting analysis of PHD3 interaction with the BC domain of ACC2. In vitro binding assay of PHD3 interaction with the BCCP domain (G) or CT domain (H). (I) ACC2 N-terminus domain interacts with PHD3. ACC2 NT-Flag and/or PHD3-HA were transfected into HEK293T cells. Proteins in cell lysates were precipitated by anti-Flag antibody and the precipitates were immunoblotted using the indicated antibodies. (J) ACC2 NT-Flag was transfected into HEK293T cells. Cells were incubated under high or low nutrient media for 8 hr. Proteins in cell lysates were precipitated by anti-Flag antibody and the precipitates were immunoblotted using hydroxyl proline antibodies.

**Figure S4.**
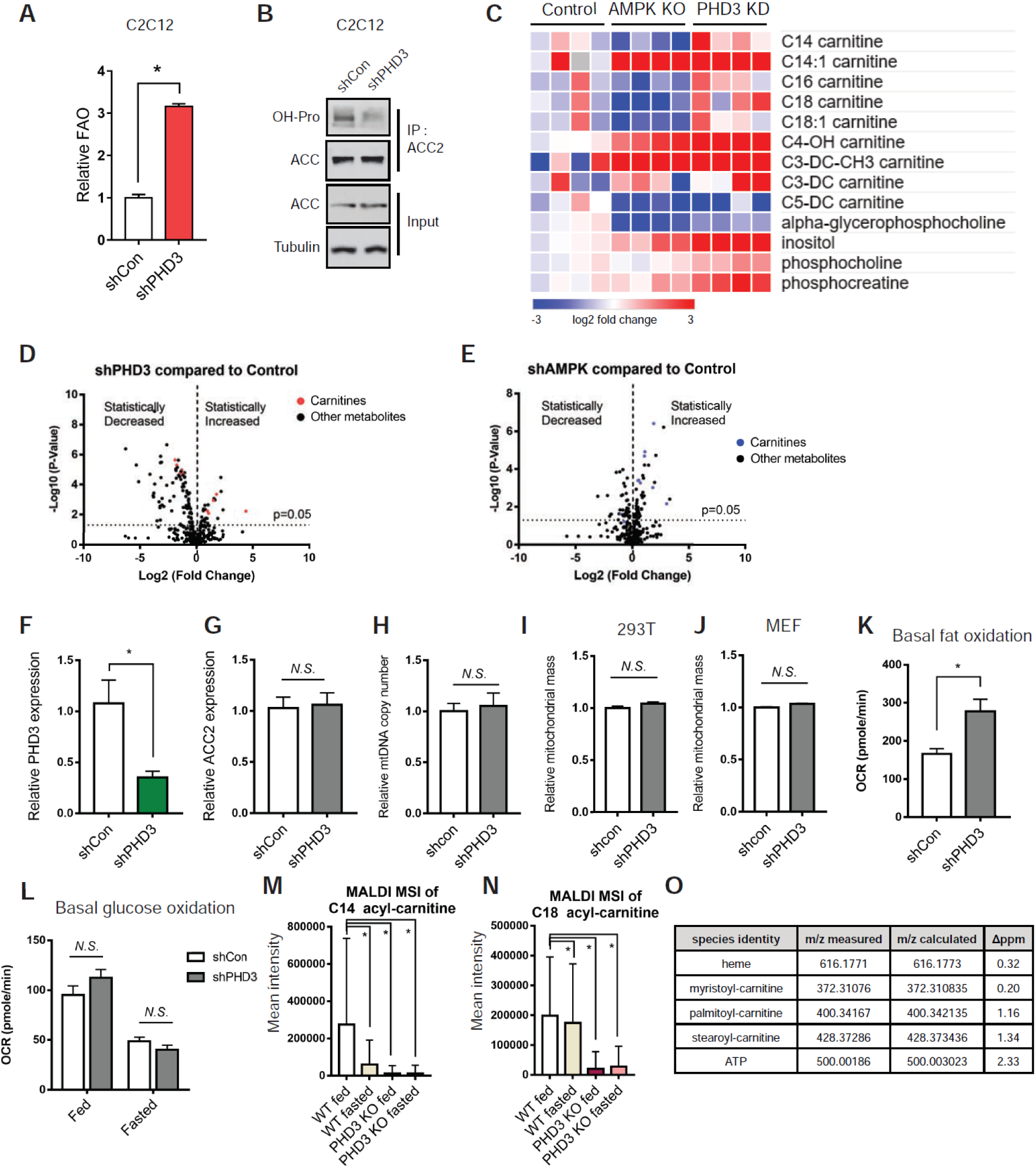
PHD3 and AMPK have differential functions in fat metabolism, but do not affect mitochondrial biogenesis. (A) Palmitate oxidation in shcontrol or shPHD3 C2C12 cells (n=3). (B) Immunoblots of anti-ACC2 immunoprecipates and corresponding whole-cell lysates of shcontrol or shPHD3 C2C12 cells. (C) Heat map depicting metabolite abundance in PHD3 knock-down, AMPKα knock out or non-silencing control MEFs. Volcano plots demonstrating that PHD3 (D) and AMPK depleted MEFs (E) have numerous statistically significant metabolic changes compared to control cells. Red and blue dots indicate each acyl-carnitine. PHD3 expression (F) or ACC2 expression (G) using B2M as a reference gene in shControl or shPHD3 HEK293T cells. (H) Mitochondrial DNA number was not changed by PHD3 knockdown. mtDNA levels were measured by MT-CO_2_ using human b-actin as a reference gene in shControl or shPHD3 HEK293T cells. Mitochondrial mass of 293T cells (I) or MEFs (J) depleted of PHD3 or control shRNA using MitoTracker Green (n = 5). (K) The basal extracellular acidification rate (ECAR) in shControl or shPHD3 MEFs (n = 15). (L) Basal oxygen consumption rate (OCR) tracing for high or low glucose cultured shControl or shPHD3 MEFs, measured using a Seahorse Bioscience XF24 Extracellular Flux Analyzer (n = 15 wells). Matrix-assisted laser desorption/ionization mass spectrometry imaging (MALDI-MSI) of myristoyl-carnitine (m/z 372.310835, Δppm = 0.2) (M), and stearoyl-carnitine (m/z 428.37286, Δppm = 1.34) (N) in WT or PHD3 KO mouse quadriceps under fed or fasted conditions. (O) Measured and calculated mass of species detected by MALDI MS (FT-ICR MS so <2 ppm expected).

**Figure S5.**
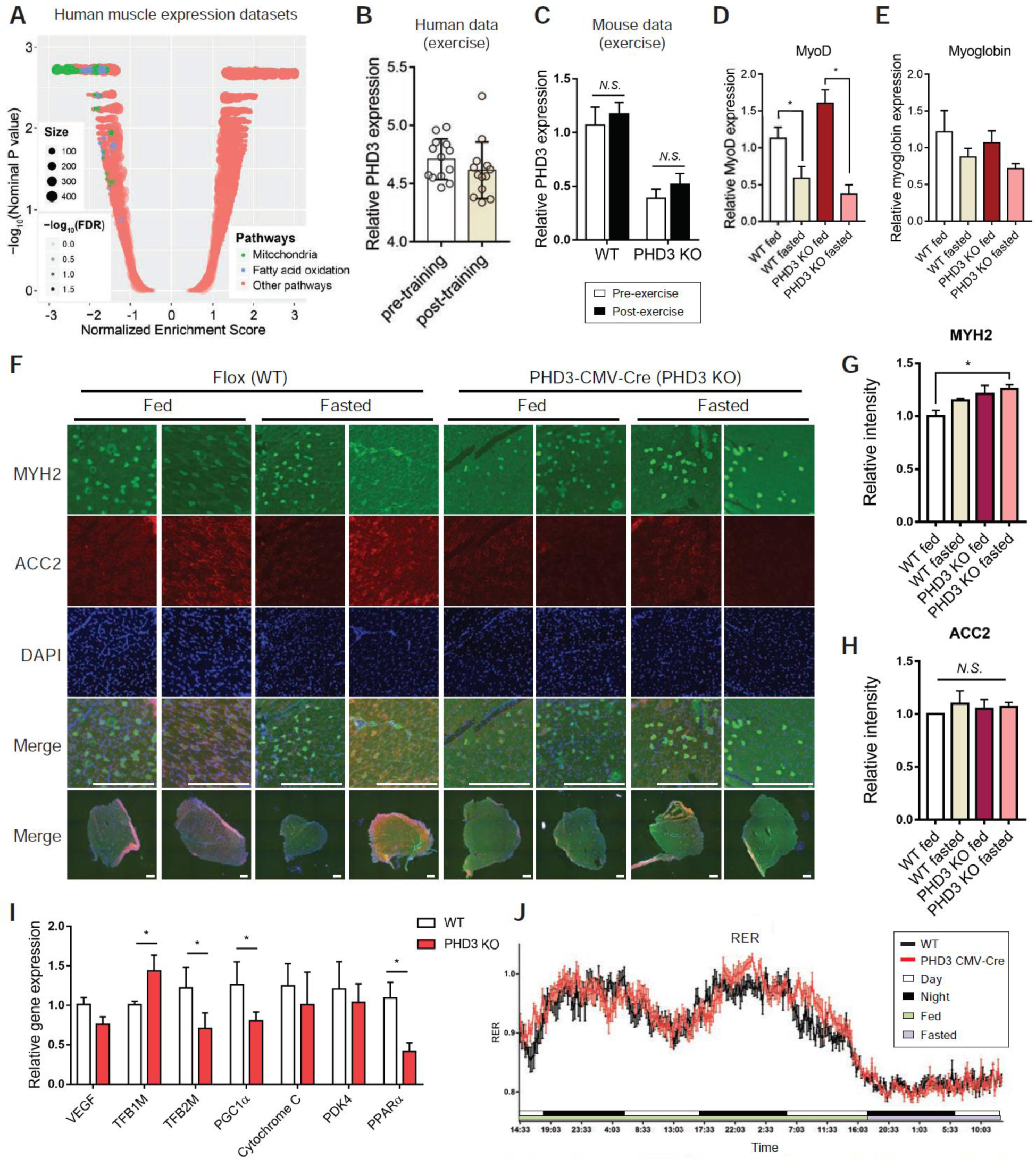
The expression of ACC2 or muscle markers were not changed under fasting conditions and PHD3 KO mouse muscle compared to WT mouse under fed conditions. (A) Gene set enrichment analysis of PHD3 indicating its negative connection with fatty acid oxidation and mitochondrial pathways in human muscle dataset (GSE3307). Mitochondrial and fatty acid oxidation pathways are highlighted in green and blue, respectively. The size of dot represents the number of genes in the pathway, and transparency indicates the significance (−log10 (FDR)) of the enrichment between this pathway and PHD3 expression. (B) PHD3 expression levels in human muscle biopsies after untraining or endurance exercise training (GSE111551). Data represent means ± SEM. (C) Relative PHD3 expression using mouse beta-actin as a reference gene in WT or PHD3 KO mice quadriceps with exercise (n=9). The mRNA levels of MyoD (D) and myoblobin (E) in WT or PHD3 KO mouse quadriceps under fed or fasting conditions (n = 4). * p < 0.05. (F) Immunofluorescence by MYH2, which is Myosin Heavy Chain Type IIA (green), ACC2 (red) and DAPI (blue) in WT or PHD3 KO mouse quadriceps under fed and fasted conditions. Scale is 500 μm. Relative intensity of MYH2 (G) and ACC2 (H). Normalized by relative intensity of DAPI. (I) Gene expression using mouse β-actin as a reference gene in WT or PHD3 KO mouse quadriceps (n = 4). (J) Respiratory exchange ratio (RER) in 20 weeks of WT or PHD3 KO mice using Comprehensive Lab Animal Monitoring System (CLAMS) (n=9). WT or PHD3 KO mice were under fed condition 48 hours and fasted condition 16 hours.

**Figure S6.**
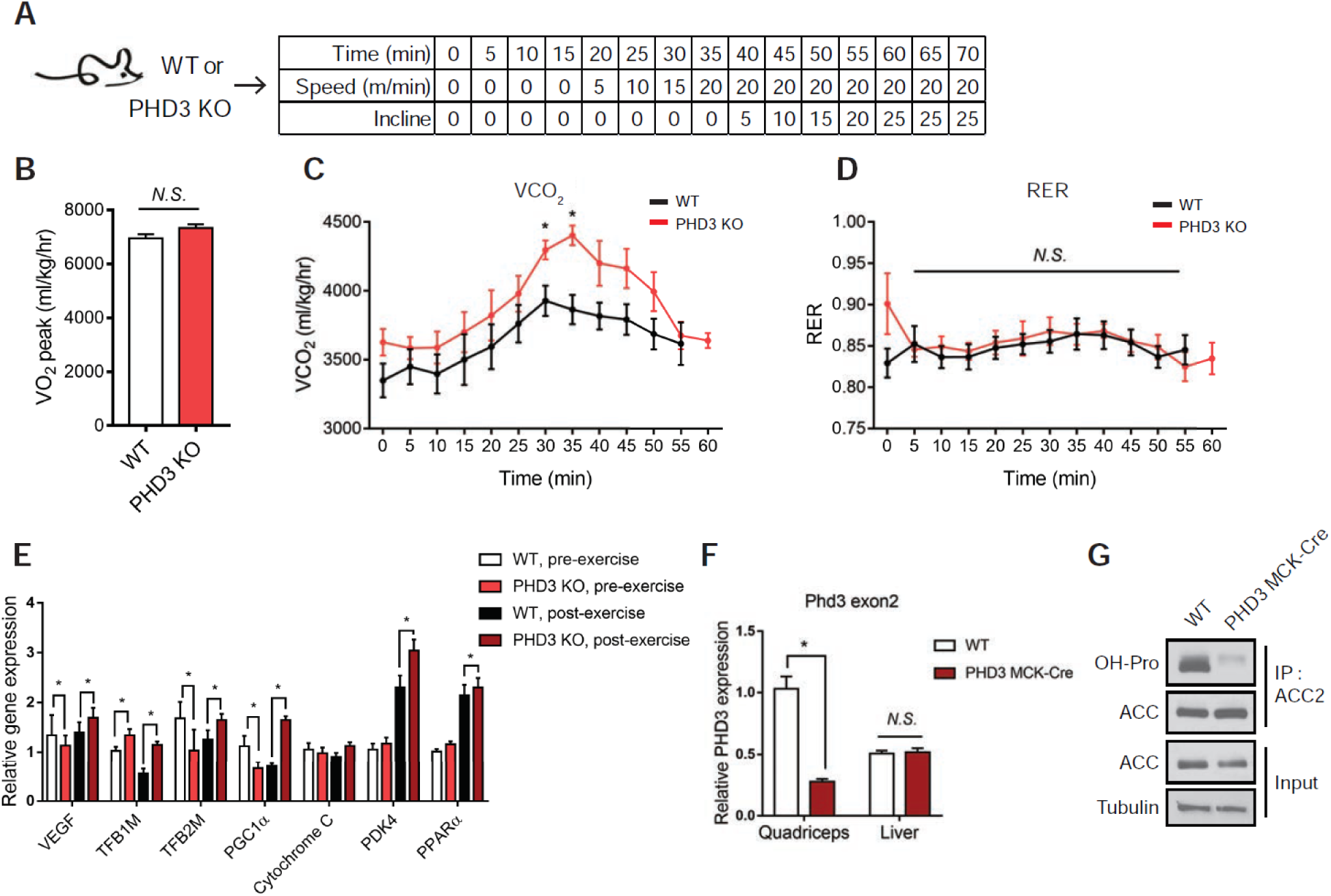
PHD3 KO mice have increased exercise capacity compared to WT. (A) Schematic of the exercise capacity treadmill experiment. (B) Age-matched mice (approx. 20 weeks old) were individually placed in metabolic cages and allowed to acclimatize for 48 h before readings were taken (n = 8 mice per group). Mean whole-body maximum oxygen consumption rate of both genotypes were calculated from the individual performances. Data represented mean ± SEM. *P < 0.05. (C) Mean whole-body CO_2_ during exercise challenge. (D) RER (VCO_2_/VO_2_). Data represented means ± SEM. *P < 0.05. (E) Gene expression using mouse b-actin as a reference gene in WT or PHD3 KO mouse quadriceps under pre- or post-exercise condition (n=4). (F) qPCR analyses to confirm the lack of WT mRNA transcripts in WT or PHD3 MCK-Cre KO mouse quadriceps and liver RNA samples. (G) Immunoprecipitation and western blotting analysis of ACC2 hydroxylation in WT or PHD3 MCK-Cre KO mice quadriceps.

**Supplement Table 1.**
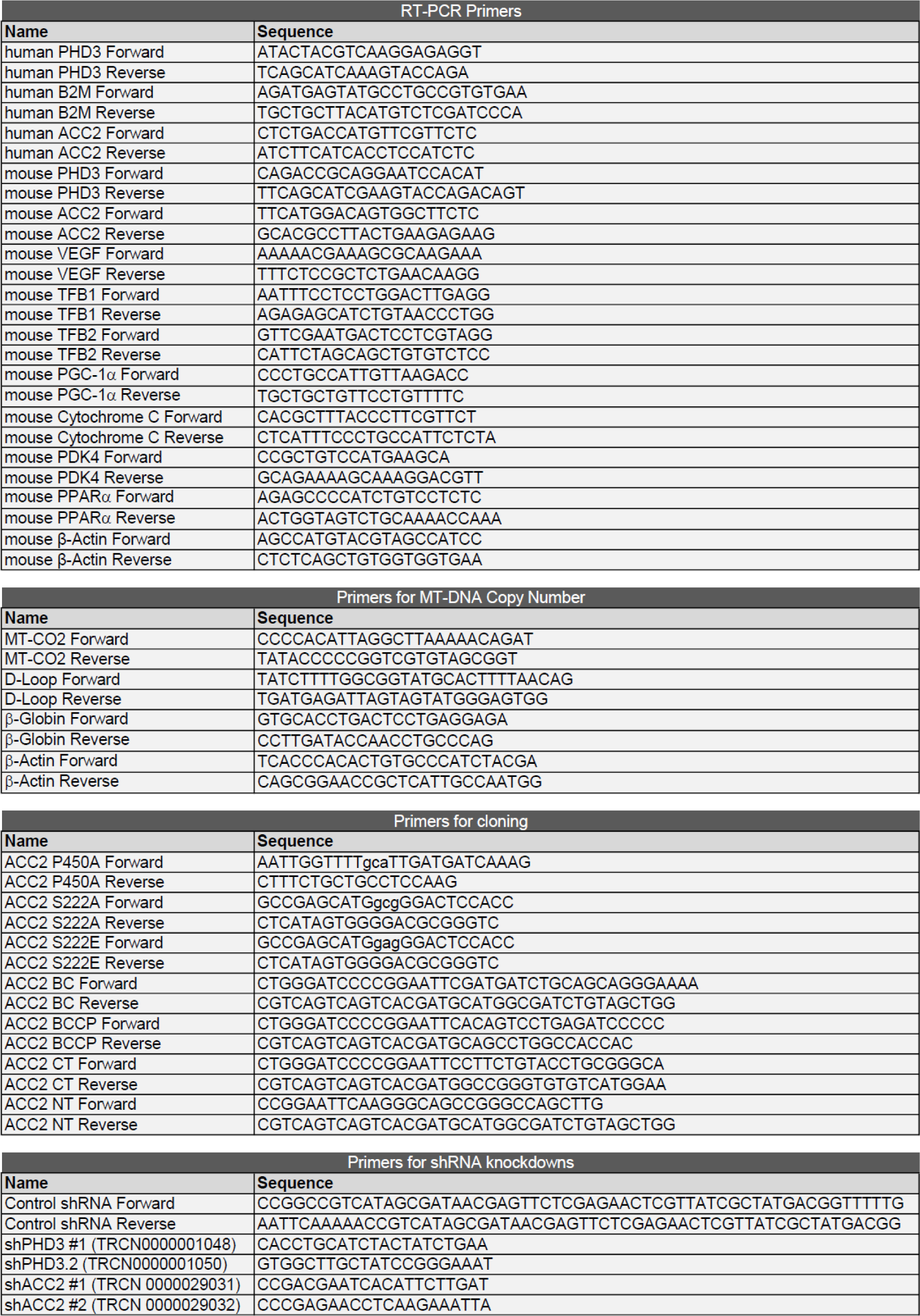
primer sequences

## REFERENCES

Abu-Elheiga, L., Oh, W., Kordari, P., and Wakil, S.J. (2003). Acetyl-CoA carboxylase 2 mutant mice are protected against obesity and diabetes induced by high-fat/high-carbohydrate diets. Proc. Natl. Acad. Sci. USA 100, 10207–10212.

Bakay, M., Wang, Z., Melcon, G., Schiltz, L., Xuan, J., Zhao, P., Sartorelli, V., Seo, J., Pegoraro, E., Angelini, C., et al. (2006). Nuclear envelope dystrophies show a transcriptional fingerprint suggesting disruption of Rb-MyoD pathways in muscle regeneration. Brain 129, 996–1013.

Berberich, M.J., Paulo, J.A., and Everley, R.A. (2018). MS3-IDQ: Utilizing MS3 Spectra beyond Quantification Yields Increased Coverage of the Phosphoproteome in Isobaric Tag Experiments. J. Proteome Res. 17, 1741–1747.

Bloemberg, D., and Quadrilatero, J. (2012). Rapid determination of myosin heavy chain expression in rat, mouse, and human skeletal muscle using multicolor immunofluorescence analysis. PLoS One. 7, e35273.

Carling, D., Clarke, P.R., Zammit, V.A., and Hardie, D.G. (1989). Purification and characterization of the AMP-activated protein kinase. Copurification of acetyl-CoA carboxylase kinase and 3-hydroxy-3-methylglutaryl-CoA reductase kinase activities. Eur. J. Biochem. 186, 129–136.

Chen, Y., Zhang, H.S., Fong, G.H., Xi, Q.L., Wu, G.H., Bai, C.G., Ling, Z.Q., Fan, L., Xu, Y.M., Qin, Y.Q., et al. (2015). PHD3 Stabilizes the Tight Junction Protein Occludin and Protects Intestinal Epithelial Barrier Function. J. Biol. Chem. 290, 20580–20589.

Cho, Y.S., Lee, J.I., Shin, D., Kim, H.T., Jung, H.Y., Lee, T.G., Kang, L.W., Ahn, Y.J., Cho, H.S., and Heo, Y.S. (2010). Molecular mechanism for the regulation of human ACC2 through phosphorylation by AMPK. Biochem. Biophys. Res. Commun. 391, 187–192.

Choi, C.S., Savage, D.B., Abu-Elheiga, L., Liu, Z.X., Kim, S., Kulkarni, A., Distefano, A., Hwang, Y.J., Reznick, R.M., Codella, R., et al. (2007). Continuous fat oxidation in acetyl-CoA carboxylase 2 knockout mice increases total energy expenditure, reduces fat mass, and improves insulin sensitivity. Proc. Natl. Acad. Sci. USA 104, 16480–16485.

Chouchani, E.T., Kazak, L., Jedrychowski, M.P., Lu, G.Z., Erickson, B.K., Szpyt, J., Pierce, K.A., Laznik-Bogoslavski, D., Vetrivelan, R., Clish, C.B., et al. (2016). Mitochondrial ROS regulate thermogenic energy expenditure and sulfenylation of UCP1. Nature 532, 112–116.

Dadgar, S., Wang, Z., Johnston, H., Kesari, A., Nagaraju, K., Chen, Y.W., Hill, D.A., Partridge, T.A., Giri, M., Freishtat, R.J., et al. (2014). Asynchronous remodeling is a driver of failed regeneration in Duchenne muscular dystrophy. J. Cell Biol. 207, 139–158.

De Leiris, J., Opie, L.H., and Lubbe, W.F. (1975). Effects of free fatty acid and enzyme release in experimental glucose on myocardial infarction. Nature 253, 746–747.

Du, W., Zhang, L., Brett-Morris, A., Aguila, B., Kerner, J., Hoppel, C.L., Puchowicz, M., Serra, D., Herrero, L., Rini, B.I., et al. (2017). HIF drives lipid deposition and cancer in ccRCC via repression of fatty acid metabolism. Nat. Commun. 8, 1769.

Efeyan, A., Comb, W.C., and Sabatini, D.M. (2015). Nutrient-sensing mechanisms and pathways. Nature 517, 302–310.

Epstein, A.C., Gleadle, J.M., McNeill, L.A., Hewitson, K.S., O’Rourke, J., Mole, D.R., Mukherji, M., Metzen, E., Wilson, M.I., Dhanda, A., et al. (2001). C. elegans EGL-9 and mammalian homologs define a family of dioxygenases that regulate HIF by prolyl hydroxylation. Cell. 107, 43–54.

Errichiello, E., Balsamo, A., Cerni, M., and Venesio, T. (2015). Mitochondrial variants in MT-CO2 and D-loop instability are involved in MUTYH-associated polyposis. J. Mol. Med. (Berl). 93, 1271–1281.

Galgani, J., and Ravussin, E. (2008). Energy metabolism, fuel selection and body weight regulation. Int. J. Obes. (Lond). *Suppl 7*, S109–119.

Galic, S., Loh, K., Murray-Segal, L., Steinberg, G.R., Andrews, Z.B., and Kemp, B.E. (2018). AMPK signaling to acetyl-CoA carboxylase is required for fasting- and cold-induced appetite but not thermogenesis. Elife 13, pii: e32656.

German, N.J., Yoon, H., Yusuf, R.Z., Murphy, J.P., Finley, L.W., Laurent, G., Haas, W., Satterstrom, F.K., Guarnerio, J., Zaganjor, E., et al. (2016). PHD3 Loss in Cancer Enables Metabolic Reliance on Fatty Acid Oxidation via Deactivation of ACC2. Mol. Cell 63, 1006–1020.

Gordon, P.M., Liu, D., Sartor, M.A., IglayReger, H.B., Pistilli, E.E., Gutmann, L., Nader, G.A., and Hoffman, E.P. (1985). Resistance exercise training influences skeletal muscle immune activation: a microarray analysis. J. Appl. Physiol. 112, 443–453.

Gowans, G.J., and Hardie, D.G. (2014). AMPK: a cellular energy sensor primarily regulated by AMP. Biochem. Soc. Trans. 42, 71–75.

Greenberg, S.A., Bradshaw, E.M., Pinkus, J.L., Pinkus, G.S., Burleson, T., Due, B., Bregoli, L., O’Connor, K.C., and Amato, A.A. (2005). Plasma cells in muscle in inclusion body myositis and polymyositis. Neurology 65, 1782–1787.

Grunt, T.W. (2018). Interacting Cancer Machineries: Cell Signaling, Lipid Metabolism, and Epigenetics. Trends. Endocrinol. Metab. 29, 86–98.

Gwinn, D.M., Shackelford, D.B., Egan, D.F., Mihaylova, M.M., Mery, A., Vasquez, D.S., Turk, B.E., and Shaw, R.J. (2008). AMPK phosphorylation of raptor mediates a metabolic checkpoint. Mol. Cell 30, 214–226.

Hardie, D.G., Ross, F.A., and Hawley, S.A. (2012). AMPK: a nutrient and energy sensor that maintains energy homeostasis. Nat. Rev. Mol. Cell Biol. 13, 251–262.

Herzig, S., and Shaw, R.J. (2018). AMPK: guardian of metabolism and mitochondrial homeostasis. Nat. Rev. Mol. Cell Biol. 19, 121–135.

Huang, Li, T., Li, X., Zhang, L., Sun, L., He, X., Zhong, X., Jia, D., Song, L., Semenza, G.L., et al. (2014). HIF-1-mediated suppression of acyl-CoA dehydrogenases and fatty acid oxidation is critical for cancer progression. Cell Rep. 8, 1930–1942.

Hunter, R.W., Treebak, J.T., Wojtaszewski, J.F., and Sakamoto, K. (2011). Molecular mechanism by which AMP-activated protein kinase activation promotes glycogen accumulation in muscle. Diabetes 60, 766–774.

Inoki, K., Zhu, T., and Guan, K.L. (2003). TSC2 mediates cellular energy response to control cell growth and survival. Cell 26, 577–590.

Jacobs, R.A., Díaz, V., Meinild, A.K., Gassmann, M., and Lundby, C. (2013). The C57Bl/6 mouse serves as a suitable model of human skeletal muscle mitochondrial function. Exp. Physiol. 98, 908–921.

Jenkins, C.M., Yang, J., Sims, H.F., and Gross, R.W. (2011). Reversible high affinity inhibition of phosphofructokinase-1 by acyl-CoA: a mechanism integrating glycolytic flux with lipid metabolism. J. Biol. Chem. 286, 11937–11950.

Laderoute, K.R., Amin, K., Calaoagan, J.M., Knapp, M., Le, T., Orduna, J., Foretz, M., and Viollet, B. (2006). 5’-AMP-activated protein kinase (AMPK) is induced by low-oxygen and glucose deprivation conditions found in solid-tumor microenvironments. Mol. Cell Biol. 26, 5336–5347.

Lee, S., Nakamura, E., Yang, H., Wei, W., Linggi, M.S., Sajan, M.P., Farese, R.V., Freeman, R.S., Carter, B.D., Kaelin, W.G. Jr., et al. (2005). Neuronal apoptosis linked to EglN3 prolyl hydroxylase and familial pheochromocytoma genes: developmental culling and cancer. Cancer Cell 8, 155–167.

Lee, J., Choi, J., Scafidi, S., and Wolfgang, M.J. (2016). Hepatic Fatty Acid Oxidation Restrains Systemic Catabolism during Starvation. Cell Rep. 16, 201–212.

Liu, X., Ide, J.L., Norton, I., Marchionni, M.A., Ebling, M.C., Wang, L.Y., Davis, E., Sauvageot, C.M., Kesari, S., Kellersberger, K.A., et al. (2013). Molecular imaging of drug transit through the blood-brain barrier with MALDI mass spectrometry imaging. Sci. Rep. 3, 2859.

Liu, Y., Ma, Z., Zhao, C., Wang, Y., Wu, G., Xiao, J., McClain, C.J., Li, X., and Feng, W. (2014). HIF-1α and HIF-2α are critically involved in hypoxia-induced lipid accumulation in hepatocytes through reducing PGC-1α-mediated fatty acid β-oxidation. Toxicol. Lett. 226, 117–123.

Manning, B.D., Toker, A. (2017). AKT/PKB Signaling: Navigating the Network. Cell 169, 381–405.

McCoin, C.S., Knotts, T.A., and Adams, S.H. (2015). Acylcarnitines--old actors auditioning for new roles in metabolic physiology. Nat. Rev. Endocrinol. 11, 617–625.

Mihaylova, M.M., and Shaw, R.J. (2011). The AMPK signalling pathway coordinates cell growth, autophagy and metabolism. Nat. Cell Biol. 13, 1016–1023.

Millay, D.P., O’Rourke, J.R., Sutherland, L.B., Bezprozvannaya, S., Shelton, J.M., Bassel-Duby, R., Olson, E.N. (2013). Myomaker is a membrane activator of myoblast fusion and muscle formation. Nature 499, 301–330.

Mul, J.D., Stanford, K.I., Hirshman, M.F., and Goodyear, L.J. (2015). Exercise and Regulation of Carbohydrate Metabolism. Prog. Mol. Biol. Transl. Sci. 135, 17–37.

Narkar, V.A., Downes, M., Yu, R.T., Embler, E., Wang, Y.X., Banayo, E., Mihaylova, M.M., Nelson, M.C., Zou, Y., Juguilon, H., et al. (2008). AMPK and PPARdelta agonists are exercise mimetics. Cell 134, 405–415.

Navarrete-Perea, J., Yu, Q., Gygi, S.P., and Paulo, J.A. (2018). Streamlined Tandem Mass Tag (SL-TMT) Protocol: An Efficient Strategy for Quantitative (Phospho)proteome Profiling Using Tandem Mass Tag-Synchronous Precursor Selection-MS3. J. Proteome Res. 17, 2226–2236.

O’Neill, H.M., Maarbjerg, S.J., Crane, J.D., Jeppesen, J., Jørgensen, S.B., Schertzer, J.D., Shyroka, O., Kiens, B., van Denderen, B.J., Tarnopolsky, M.A., et al. (2011). AMP-activated protein kinase (AMPK) beta1beta2 muscle null mice reveal an essential role for AMPK in maintaining mitochondrial content and glucose uptake during exercise. Proc. Natl. Acad. Sci. USA 108, 16092–16097.

O’Neill, H.M., Lally, J.S., Galic, S., Thomas, M., Azizi, P.D., Fullerton, M.D., Smith, B.K., Pulinilkunnil, T., Chen, Z., Samaan, M.C., et al. (2014). AMPK phosphorylation of ACC2 is required for skeletal muscle fatty acid oxidation and insulin sensitivity in mice. Diabetologia 57,1693–1702.

Palm, W., and Thompson, C.B. (2017). Nutrient acquisition strategies of mammalian cells. Nature 546, 234–242.

Paulo, J.A. (2016). Sample preparation for proteomic analysis using a GeLC-MS/MS strategy. J. Biol. Methods 3, pii: e45.

Randle, P.J., Garland, P.B., Hales, C.N., and Newsholeme, E.A. (1963). The glucose fatty-acid cycle. Its role in insulin sensitivity and the metabolic disturbances of diabetes mellitus. Lancet 1, 785–789.

Raue, U., Trappe, T.A., Estrem, S.T., Qian, H.R., Helvering, L.M., Smith, R.C., and Trappe, S. (1985). Transcriptome signature of resistance exercise adaptations: mixed muscle and fiber type specific profiles in young and old adults. J. Appl. Physiol. 112, 1625–1636.

Ruzankina, Y., Pinzon-Guzman, C., Asare, A., Ong, T., Pontano, L., Cotsarelis, G., Zediak, V.P., Velez, M., Bhandoola, A., and Brown, E.J. (2007). Deletion of the developmentally essential gene ATR in adult mice leads to age-related phenotypes and stem cell loss. Cell Stem Cell 1, 113–126.

Salt, I.P., Johnson, G., Ashcroft, S.J., and Hardie, D.G. (1998). AMP-activated protein kinase is activated by low glucose in cell lines derived from pancreatic beta cells, and may regulate insulin release. Biochem. J. 335, 533–539.

Spinelli, J.B., and Haigis, M.C. (2018). The multifaceted contributions of mitochondria to cellular metabolism. Nat. Cell Biol. 20, 745–754.

Spriet, L.L. (2014). New insights into the interaction of carbohydrate and fat metabolism during exercise. Sports Med. *Suppl 1*, S87–96.

Takeda, K., Ho, V.C., Takeda, H., Duan, L.J., Nagy, A., and Fong, G.H. (2006). Placental but not heart defects are associated with elevated hypoxia-inducible factor alpha levels in mice lacking prolyl hydroxylase domain protein 2. Mol. Cell Biol. 26, 8336–8346.

Tennant, D.A., and Gottlieb, E. (2010). HIF prolyl hydroxylase-3 mediates alpha-ketoglutarate-induced apoptosis and tumor suppression. J. Mol. Med. (Berl). 88, 839–849.

Tong, L., and Harwood, H.J. Jr. (2006). Acetyl-coenzyme A carboxylases: versatile targets for drug discovery. J. Cell Biochem. 99, 1476–1488.

Tran, T.H., Hsiao, Y.S., Jo, J., Chou, C.Y., Dietrich, L.E., Walz, T., and Tong, L. (2015). Structure and function of a single-chain, multi-domain long-chain acyl-CoA carboxylase. Nature 518, 120–124.

Wei, J., and Tong, L. (2015). Crystal structure of the 500-kDa yeast acetyl-CoA carboxylase holoenzyme dimer. Nature 526, 723–727.

Woods, S.C., and Ramsay, D.S. (2011). Food intake, metabolism and homeostasis. Physiol. Behav. 104, 4–7.

